# The largest SWI/SNF polyglutamine domain is a pH sensor

**DOI:** 10.1101/165043

**Authors:** J. Ignacio Gutiérrez, Greg Brittingham, Xuya Wang, David Fenyö, Liam J. Holt

## Abstract

Polyglutamines are known to form aggregates in pathogenic contexts, such as in Huntington’s disease, however little is known about their role in normal biological processes. We found that a polyglutamine domain in the SNF5 subunit of the yeast SWI/SNF complex, histidines within this sequence, and transient intracellular acidification are required for efficient transcriptional regulation during carbon starvation. We hypothesized that a pH-driven oligomerization of the SNF5 polyglutamine region is required for transcriptional reprogramming. In support of this idea, we found that a synthetic spidroin domain from spider silk, which is soluble at pH 7 but oligomerizes at pH ~ 6.3, could partially complement the function of the SNF5 polyglutamine. These results suggest that the SNF5 polyglutamine domain acts as a pH-driven transcriptional regulator.

## Introduction

Intracellular pH (pH_i_) influences all biological processes by determining the protonation state of biological molecules, including the charged amino acid side-chains of proteins, and particularly histidines, which have a near-neutral pKa (Whitten, Garcia-Moreno E. and Hilser, 2005). Early work suggested that pH_i_ was constant during development (Needham, 1926). However, more advanced technologies have since revealed that pH_i_ does vary, and that changes in pH_i_ can regulate metabolism (Busa and Nuccitelli, 1984; Young *et al*., 2010), proliferation (Busa and Crowe, 1983), and cell fate (Okamoto, 1994), among other processes.

The budding yeast *Saccharomyces cerevisiae* is adapted to an acidic environment, and standard growth media is typically at pH 4.0 – 5.5. The plasma membrane proton pump Pma1 and the V-ATPases maintain near neutral pH_i_ by pumping protons out of the cell and to the vacuole, respectively (Martínez-Muñoz and Kane, 2008). When cells are starved for carbon, the Pma1 pump and the V-ATPases are inactivated, leading to a rapid acidification of the intracellular space to pH ~ 6 (Kane, 1995; Orij *et al*., 2009).

Acidification of intracellular pH upon carbon-starvation is thought to conserve energy, leading to storage of metabolic enzymes in filamentous aggregates (Petrovska *et al*., 2014), reduction of macromolecule diffusion (Joyner *et al*., 2016; Munder *et al*., 2016), decreased membrane biogenesis (Young *et al*., 2010) and possibly a partial phase transition of the cytoplasm into a solid-like structure (Joyner *et al*., 2016; Munder *et al*., 2016). These studies demonstrate that many physiological processes are inactivated when pH_i_ drops, an adaptation, which may aid the cell in surviving stress. However, numerous processes must also be upregulated during carbon starvation to enable adaptation to this stress. In particular, the cell must induce expression of glucose-repressed genes (DeRisi, 1997; Zid and O’Shea, 2014). However, in the current literature there is no evidence of a relationship between an acidic pH_i_ and stress-gene induction.

The Sucrose Non Fermenting genes (SNF) were among the first genes found to be required for induction of glucose-repressed genes (Neigeborn and Carlson, 1984). Several of these genes were later identified as members of the SWI/SNF complex (Abrams, Neigeborn and Carlson, 1986; Carlson, 1987), an 11 subunit chromatin remodeling complex that is highly conserved from yeast to mammals (Peterson and Herskowitz, 1992; Chiba *et al*., 1994; Peterson, Dingwall and Scott, 1994). The SWI/SNF complex affects the expression of ~10% of the genes in *Saccharomyces cerevisiae* during vegetative growth (Sudarsanam *et al*., 2000). Upon carbon starvation, most genes are down-regulated, but a set of glucose-repressed genes needed for adaptation to nutritional stress are strongly induced (Zid and O’Shea, 2014). The SWI/SNF complex is required for the efficient expression of several hundred stress-response and glucose-repressed genes (Sudarsanam *et al*., 2000; Biddick *et al*., 2008). Sequence analysis reveals a strong enrichment of low-complexity sequence in the complex. In particular, 4/11 subunits contain polyglutamines (polyQ) or glutamine-rich low complexity sequence (LCS).

PolyQs have been predominantly studied in the disease context. Nine neurodegenerative illnesses, including Huntington’s disease, are thought to be caused by neurotoxic aggregation seeded by proteins that contain expanded polyQs (Fan *et al*., 2014). However, polyQs are relatively abundant in *Eukaryotic* cells: More than 100 human proteins contain polyQs, and the *Dictyostelium* and *Drosophilid* phyla have polyQ structures in ~10% and ~5% of their proteins respectively (Schaefer, Wanker and Andrade-Navarro, 2012). Furthermore, there is clear evidence of purifying selection to maintain polyQs in the *Drosophilids* (Huntley and Clark, 2007). This prevalence and conservation suggests an important biological function for these repeats. Glutamine rich transactivation domains (TAD) are an example of functional polyQs (Kadonaga *et al*., 1987, 1988), however their precise role in transcription remains poorly understood. Some research suggests that glutamine-rich TADs induce transcription by binding to transcription factors (Prochasson *et al*., 2003), and recruitment of transcriptional machinery (Laurent, Treitel and Carlson, 1990; Geng, Cao and Laurent, 2001; Janody *et al*., 2001). Beyond transcription, recent work in *Ashbya gosypii* has revealed a structural role for polyQ-containing proteins through phase separation into liquid droplets to enable subcellular localization of signaling molecules (Zhang *et al*., 2015).

The *SNF5* regulatory subunit contains the largest polyQ of the SWI/SNF complex; this N-terminal LCS domain contains 121 glutamine residues. We investigated the relationship between the *SNF5* polyQ domain and the cytosolic acidification that occurs during glucose-starvation. By single cell analysis, we found that intracellular pH is highly dynamic and varies between subpopulations of cells within the same culture. After an initial decrease to pH_i_ <6.5, a subset of cells recovered their pH_i_ to ~7. This transient acidification was required for expression of glucose-repressed genes. The *SNF5* polyQ activation domain was also required for timely gene expression, including four histidines that we identify as putative pH sensors. We propose transient aggregation of low complexity sequences as a possible mechanism for transcriptional reprograming and provide support for this model using an orthogonal pH-dependent aggregation domain.

## Results

### The N-terminal polyQ domain of *SNF5* affects the viability of carbon-starved yeast cells in a pH-dependent manner

We compared the fitness of WT yeast to strains with a complete deletion of the *SNF5* gene (*snf5∆*), or with a precise deletion of the N-terminal polyQ domain of *SNF5*, referred to as *∆Q*-*snf5* (figure 1A, B). In rich media, the *snf5∆* strain has a severe growth defect (figure 1C) (Laurent, Treitel and Carlson, 1990), while the *∆Q*-*snf5* grows at a similar rate to wild-type control (figure 1C) (Prochasson *et al*., 2003). Next, we compared the viability of these strains after an acute switch to synthetic complete medium with no glucose for 24 hours. Cells were plated before and after starvation and colonies were counted to assess viability. No difference was apparent; all strains maintained 100% viability after starvation (Fig 1D - pH: 6.5). Therefore, we reasoned that we would need to increase the sensitivity of our assay to reveal a phenotype for our mutants.

**Figure 1:**
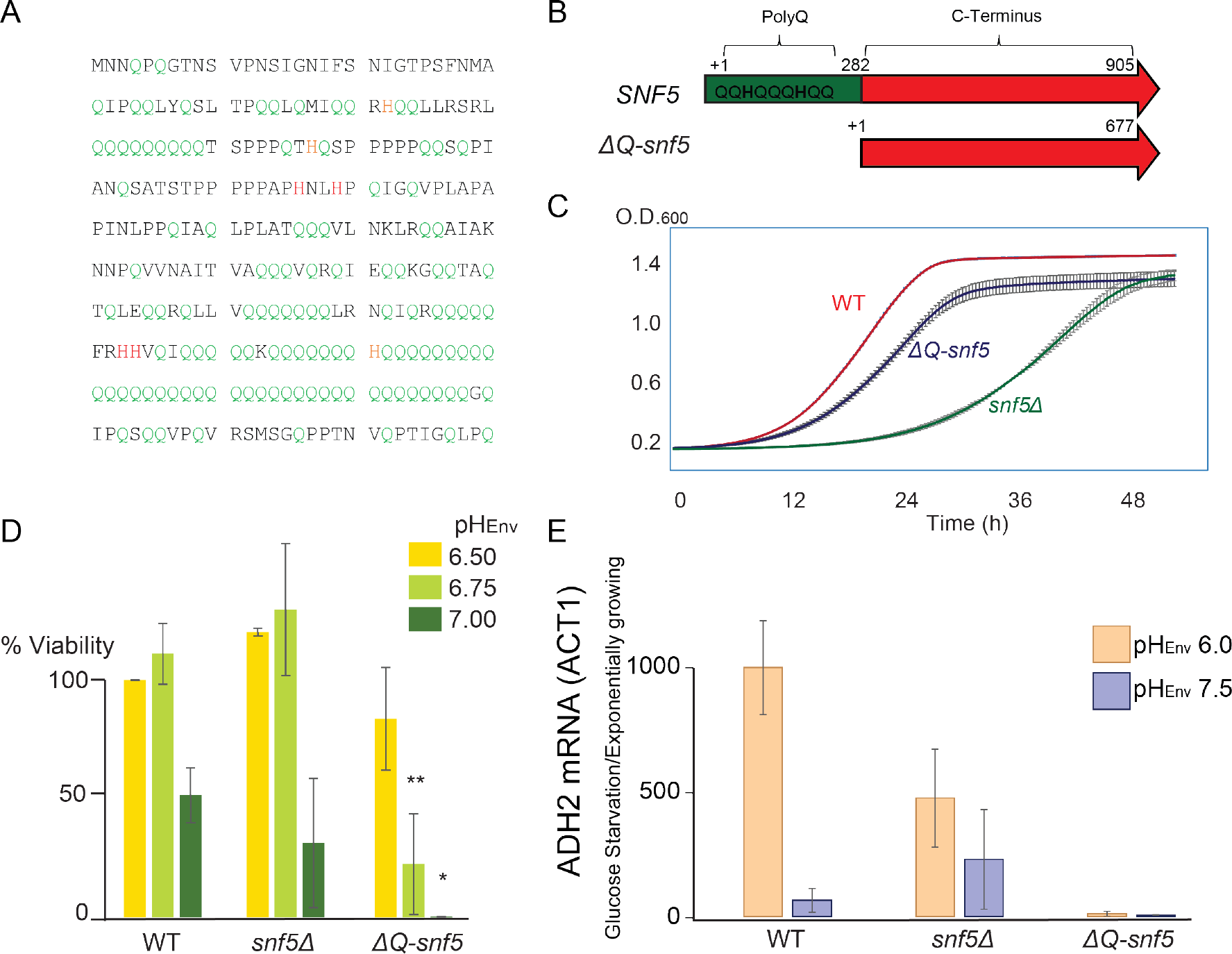
Deletion of the *SNF5* polyglutamine leads to loss of viability in carbon starvation, especially at neutral environmental pH. A- The 288 amino acids of the polyQ domain of *SNF5*, glutamine is shown in green and histidine in red/orange. B- Schematic representation of the *SNF5* gene with its polyQ domain indicated at the N-terminus (Green). C - Growth curves for WT, *ΔQ-snf5* and *snf5Δ* strains in rich media. D- Viability after 24 h of glucose-starvation at pH_env_ 6.5 (Yellow), 6.75 (Light Green) and 7.0 (Dark Green) p-values are Student’s T-test compared to WT, * indicates < 0.05 and ** indicates < 0.005. E- RT-QPCR experiments to determine *ADH2* mRNA content after 4 h of glucose-starvation at pH_env_ 6.0 (orange) and pH_env_ 7.5 (blue). *ACT1* mRNA was used as a control; y-axis corresponds to fold induction of *ADH2* mRNA on glucose-starvation over exponentially growing cells.

Intracellular acidification has been shown to be important for viability during carbon starvation (Munder *et al*., 2016). Accordingly, when we buffered the environmental pH (pH_env_) to 7, thus preventing intracellular acidification, the viability of WT cells dropped to 50% (figure 1D). Under this same condition *∆Q*-*snf5* strains completely lost viability. In contrast, the *snf5∆* strain had similar survival as WT at all pH_env_ values tested (figure 1D), although *snf5∆* colonies were smaller than those of WT and ∆*Q*-*snf5* strains, consistent with growth defects measured in liquid culture (figure 1C). Together, these results show that the *snf5∆* and *∆Q*-*snf5* alleles are different.

To ensure that deletion polyQ of *SNF5* does not alter Snf5p levels, we performed a Western blot analysis and observed similar levels of Snf5p for all alleles during glucose starvation at both pH 5.0 and 7.5 (supplemental figure 1). It has been shown by immunoprecipitation that while deletion of the whole *SNF5* gene breaks down the SWI/SNF complex into smaller sub-complexes, deletion of the N-terminal domain of Snf5p (*∆Q*-*snf5*) does not impair the architecture of the SWI/SNF complex (Prochasson *et al*., 2003) (Dutta *et al*., 2017; Sen *et al*., 2017). These results indicate that the phenotypes that we observed in the *∆Q*-*snf5* allele are not due to Snf5 protein degradation or disruption of the SWI/SNF complex.

### Induction of the SWI/SNF target *ADH2* in response to carbon starvation requires an acidic environment and the polyQ domain of *SNF5*

Alcohol dehydrogenase 2 (*ADH2*) is tightly repressed in the presence of glucose, and is strongly induced upon carbon starvation (Biddick *et al*., 2008) and the SWI/SNF complex is required for the efficient and timely induction of *ADH2* (Young *et al*., 2008). Therefore, we chose *ADH2* as a model for gene induction studies. By RT-Q-PCR, we found that after 4 hours of acute carbon starvation at pH_env_ 6, wild-type cells induced *ADH2* transcription approximately 1000-fold, while *∆Q*-*snf5* did not induce detectable levels of transcript (figure 1E). Neither WT nor *∆Q*-*snf5* strains induced *ADH2* when the starvation media was adjusted to pH_env_ 7.4. In contrast the *snf5∆* strain expressed *ADH2* at lower levels than control in normal starvation, but maintained some expression when the starvation media was adjusted to pH_env_ 7.4 (figure 1E). These results suggest that the *SNF5* subunit is repressive for transcription but somehow intracellular acidification during starvation leads to derepression through a mechanism that requires the polyQ domain.

To facilitate the further study of the relationship between environmental pH and *ADH2* induction, and to enable the study of gene regulation at a single-cell level, we generated a reporter gene with the mCherry fluorescent protein under the control of the *ADH2* promoter (*P_ADH2_-mCherry*). Time course experiments showed that in optimal conditions, full transcription and translation of the mCherry protein took 24 hours, but robust induction was apparent at 6 hours (figure 2A). Induction of *P_ADH2_-mCherry* was clearly bimodal for WT cells, indicating substantial cell-to-cell variability in *ADH2* promoter activity under our conditions. This bimodality of *ADH2* induction was lost in the *snf5∆* null strain (figure 2A). These data indicate that the polyQ of *SNF5* is required for bimodal induction of *ADH2*. As expected, the phenotype of the *∆Q*-*snf5* strain was far more severe, this strain completely failed to induce *P_ADH2_-mCherry* (figure 2A). These results are consistent with previous reports where deletion of *SNF5* and *SWI1* polyQs had a more severe phenotype than the double null mutant in some conditions (Prochasson *et al*., 2003). Indeed, we found that *∆Q-snf5* acts as a dominant-negative mutation in strains bearing WT and *∆Q-snf5* alleles in the endogenous location (supplemental figure 2).

**Figure 2:**
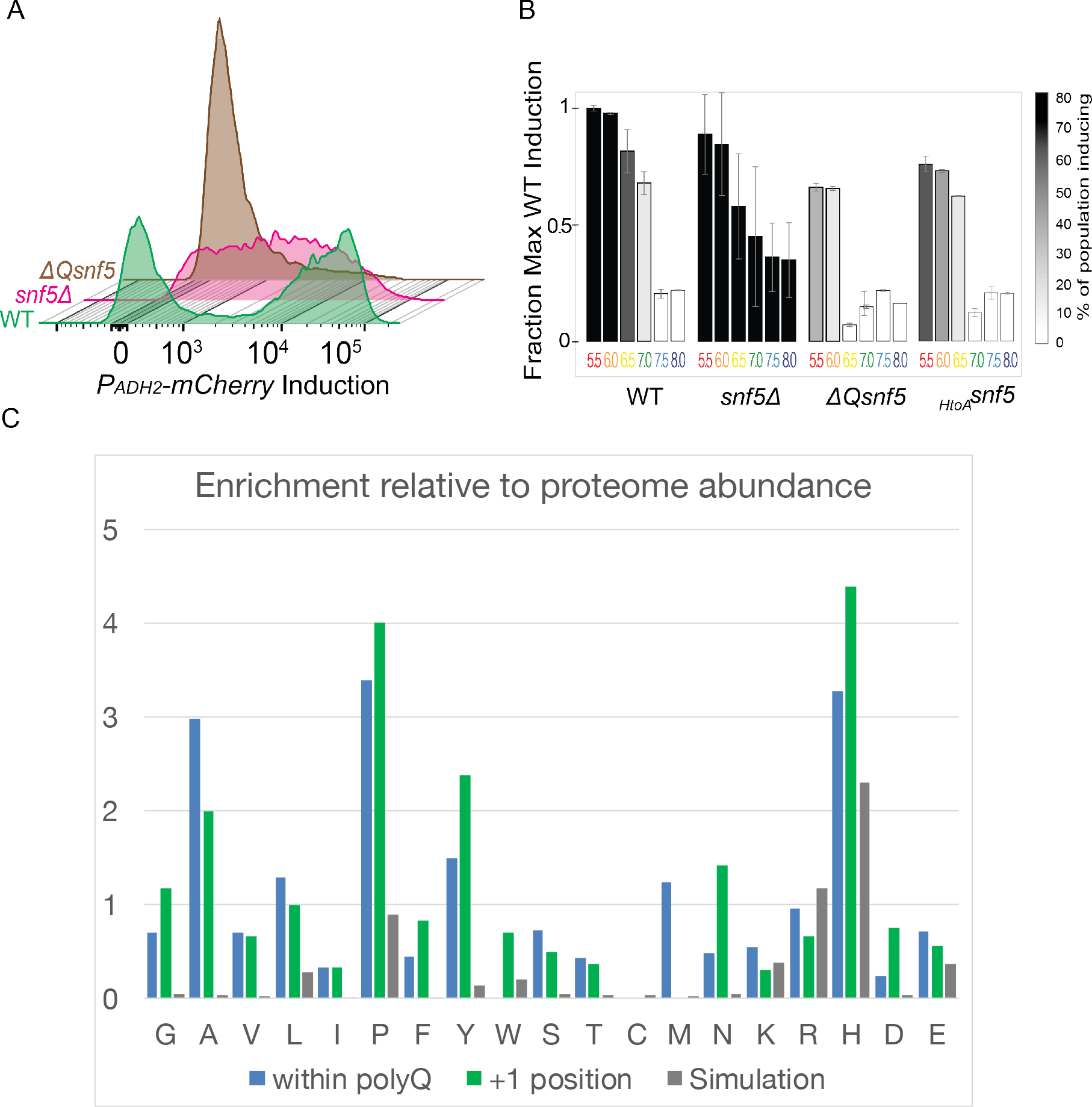
*ADH2* expression requires an acidic environmental pH and the *SNF5* polyQ. A- Example of flow cytometry histograms of *P_ADH2_-mCherry* induction after 6 h of glucose starvation, 10000 cells of each strain are plotted. B- Expression of *P_ADH2_-mCherry*, upon glucose starvation at the indicated pH_env_. Raw data was quantified by fitting to single or double Gaussian models (see material and methods) to quantify relative levels of gene induction (bar height) and the fraction of cells inducing (bar shade). C- Sequence analysis results for all *Saccharomyces cerevisiae* polyQs, showing frequencies of each amino acid compared to the proteome average, within polyQ structures (blue), and immediately C-terminal to the structures (the +1 position, green). We also show expected frequencies generated for a null-hypothesis where all sequences are generated from a pure polyQ, followed by neutral evolution – results are averages of 10,000 simulations (grey).

We next explored the effect of environmental pH on *P_ADH2_-mCherry* expression. WT cells robustly induced *P_ADH2_-mCherry* at all pH_env_ below 6.5, but almost completely failed to induce the reporter at pH_env_ of 7.0 and above (figure 2B, supplemental figure 3). In contrast the *∆Q-snf5* strain failed to induce *P_ADH2_-mCherry* at any pH_env_. The *snf5∆* strain showed a decrease of induction efficiency in control conditions (pH_env_: 5.5) as previously reported (Biddick *et al*., 2008; Biddick, Law and Young, 2008), and consistent with our observations by RT-Q-PCR, the *snf5∆* strain showed some induction at neutral pH_env_ while the wild-type completely failed to induce (figure 2B, supplemental figure 3). Taken together, these results suggest that *SNF5* is a pH-sensitive regulator of SWI/SNF function that plays both positive and negative regulatory roles. In this model, loss of the polyQ domain could lock the SWI/SNF complex in a state that is repressive for glucose repressed genes, while deletion of the whole *SNF5* gene produces derepression of the *ADH2* gene on media with neutral pH. We conclude that both the polyQ of *SNF5* and an acidic environment are required for robust regulation of *ADH2* induction upon carbon starvation.

### Histidines are enriched after polyQ sequences in multiple species

We reasoned that polyQ alone would not be a sensitive pH sensor as glutamine does not change properties over biologically relevant pH ranges. Therefore, to gain further functional insight, we analyzed amino acid enrichments within and around glutamine-rich low complexity sequences. Here, our functional definition of polyQ is a polypeptide sequence containing at least ten glutamines interrupted by no more than two non-glutamine amino acid residues. Within the *S. cerevisiae* proteome, there is a strong enrichment (>3 fold) for histidine and proline within polyQ sequences compared to the global frequency of these amino acids (figure 2C). For histidine, this enrichment is even more pronounced immediately C-terminal to polyQ repeats (>4 fold). Enrichment of histidines within and immediately after polyQs is also apparent in the human, *Drosophila melanogaster* and *Dictyostelium discoidum* proteomes (supplemental figure 4). This enrichment for certain amino acids could be an indication of functional importance, however the codons encoding glutamine (CAG and CAA) are similar to those for histidine (CAT and CAC), therefore random mutations within polyQ-encoding DNA will tend to create histidines with higher probability than other amino acids. Therefore, we tested a null hypothesis that the observed structure of polyQ sequence is due to generation of polyQ repeats (perhaps through CAG repeat expansion) and then mutation of the underlying DNA sequence in the absence of selection. We ran simulations where we allowed CAG, CAA and mixed CAGCAA repeats (artificial polyQ-encoding sequences) to randomly mutate at the nucleotide substitution frequencies that have been empirically described for *S. cerevisiae* (Zhu *et al*., 2014) until the polyQ had degenerated to have around 20% non-Q amino acids (the same frequency observed in polyQs of *S. cerevisiae*). The average results of 10,000 simulations are presented as grey bars in figure 2C. While this model does predict enrichment of histidines, the real polyQ structures are significantly more enriched suggesting that there is selective pressure to embed histidine residues within and adjacent to these structures.

### pH changes are sensed by histidines in the polyQ of *SNF5*

The imidazole side chain of histidine has a pKa of around 6. Therefore, its protonation state may serve as pH sensor. For example, the protonation state due to changes in intracellular pH of a single histidine residue determines the activation of the small G-protein RasGRP1 (Vercoulen *et al*., 2017). The N-terminal polyQ of Snf5p contains 6 histidines (shown in red and orange in figure 1A). Technical difficulties related to PCR and gene-synthesis within low complexity sequence frustrated attempts to replace all 6 histidines, however, we were able to mutate 4 of 6 histidines to alanine (shown in red in figure 1B). We refer to this allele as *_HtoA-_snf5*.

The *_HtoA-_snf5* allele phenotype was almost as severe as that of a complete deletion of the polyQ. Under optimal pH conditions (pH_env_ 5.5) both the expression level and fraction of cells that induced was reduced in *_HtoA-_snf5* strains compared to WT (figure 2B, supplemental figure 3). Under less favorable conditions of pH_env_ 6.5 where WT still strongly induced ADH2-mCherry, *_HtoA-_snf5* showed no induction at all. These results indicate that removal of histidines from the Snf5 N-terminus desensitizes the system to pH change and suggests that these histidines are important for the pH sensing function of *SNF5*.

### Single cell analysis reveals a bimodal pH response to glucose starvation, and pH recovery precedes *ADH2* expression

Intracellular pH can be studied with pHluorin, a GFP derivative that has been engineered as a ratiometric pH indicator (Miesenböck, De Angelis and Rothman, 1998). Previous studies with pHluorin reported that the cytosolic pH of *S. cerevisiae* drops to around 6 during carbon starvation (Orij *et al*., 2009). However, these were average population measurements, partly because expression levels between cells can vary when expressed from *CEN/ARS* plasmids, as they were in the original constructs. We expressed the pHluorin gene from the strong *TDH3* promoter and integrated this construct into the *URA3 locus*. Our reengineered pHluorin gave strong expression with less noise than previous systems, enabling accurate single cell pH_i_ measurements. We compared the pH_i_ of cells with either nuclear or cytoplasmic pHluorin and found that pH_i_ dynamics are the same in both compartments under glucose starvation (data not shown). We generated strains with both the *P_ADH2_-mCherry* and *PTDH3-pHluorin* reporters, to investigate the dynamics of both pH change and *ADH2* expression.

In WT cells, the pH_i_ of the entire population dropped to < 6.5 immediately after glucose was removed in starvation media pH_env_ 5.5. Then, after 30 minutes, the culture began to split into 2 subpopulations: Around half of the cells further acidified to pH_i_ 6, and did not show any *P_ADH2_-mCherry* induction, while the other half recovered their pH_i_ to ~ 7 after 2 hours and strongly induced *P_ADH2_-mCherry* after 4 hours (figure 3A, first panel, quantified in figure 3C, first panel). WT cells subjected to acute glucose-starvation at pH_env_ 7.4 mildly dropped their pH_i_, but not below 6.5 (figure 3B). Experiments with cycloheximide showed that the initial cytosolic acidification was independent of gene expression, but subsequent neutralization requires protein translation (supplemental figure 6).

**Figure 3:**
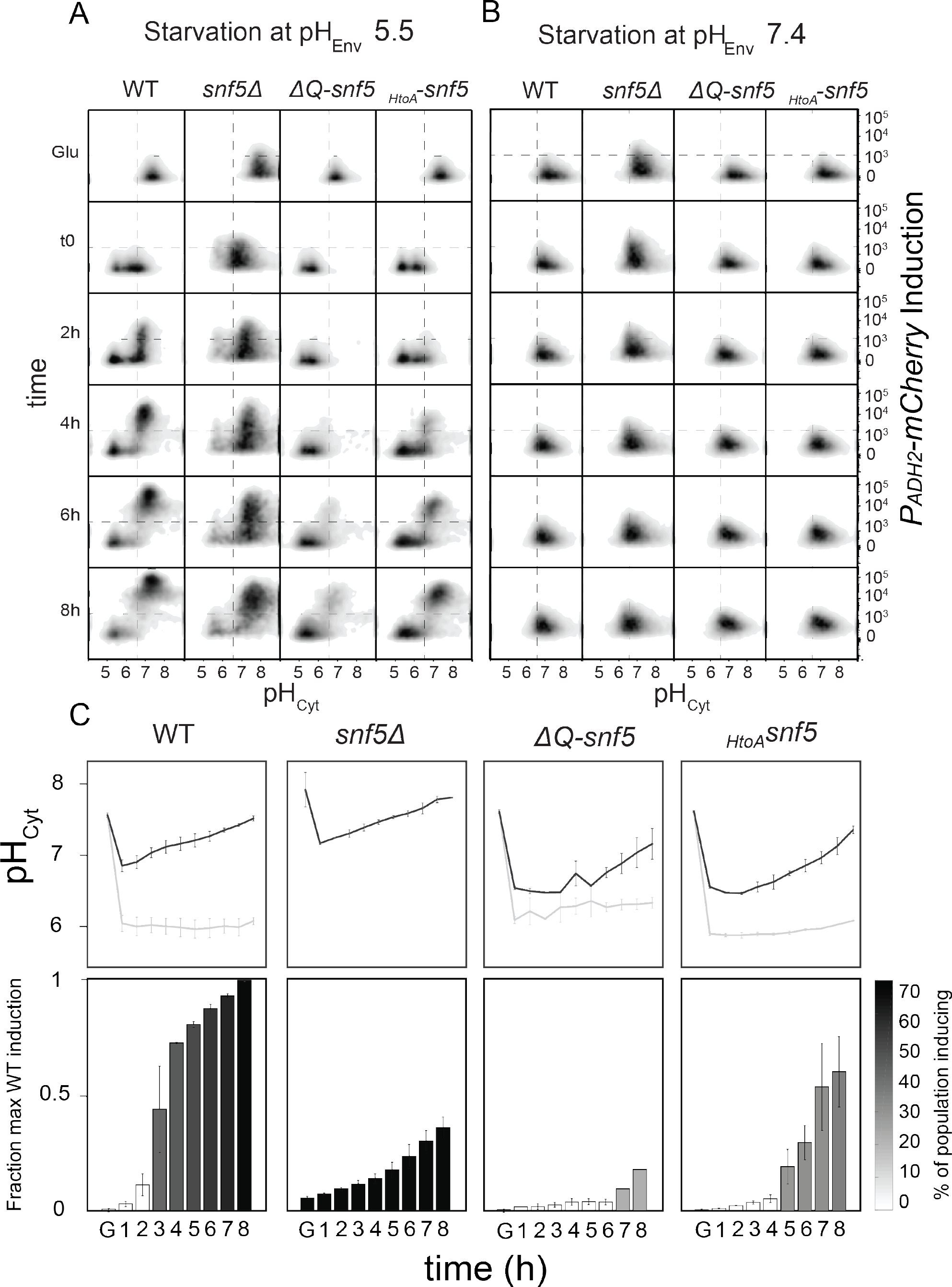
Single cell analysis reveals a bifurcation in behavior upon glucose-starvation: One subpopulation acidifies to pH 6 and fails to induce, while a second subpopulation transiently acidifies and then induces *ADH2*. A- Dot plots of cytometry data quantifying cytosolic pH (x-axis) and induction of *P_ADH2_-mCherry* (y-axis) in glucose-starvation media at pH_env_ 5.5. 10,000 cells were sampled at each time point. B- Same than A, but on glucose-starvation media pH_env_ 7.5. C- Raw data from A was fitted to single or double Gaussian models (see materials and methods). Top panels show quantification of cytosolic pH, each time point has one or two peaks and where there are two lines (black and grey), these represent the median pH values for the more neutral and more acidic populations respectively. Bottom panels show levels of induction of the *P_ADH2_-mCherry* expression. Bar height indicates to the intensity of *mCherry* fluorescence (normalized to maximum values for the WT control strain) and the color code represents the percentage of the population that induces the *P_ADH2_-mCherry* reporter above background levels.

### *SNF5* mutants have an altered pH response upon glucose starvation

We found that *SNF5* plays a role in pH regulation during acute glucose starvation. The *snf5∆* strain did not acidify as strongly as the wild-type strain, and bifurcation in pH_i_ and *ADH2* expression did not occur (figure 3A and C, second panel). In contrast, the *∆Q-snf5* mutant acidified to pH_i_ 6 but then few cells recovered to pH 7 (figure 3A and C, third panel). The *_HtoA-_snf5* mutant acted as a hypomorph – mostly phenocopying *∆Q-snf5* but a fraction of cells neutralized and induced *ADH2* at lower levels than WT cells (figure 3A and C, fourth panel).

Together, these results indicate that *SNF5* is involved in pH regulation upon glucose starvation. The C-terminus of the Snf5 protein is required for proper acidification of the cytosol, while the polyQ domain, with histidines intact, is required for the recovery to neutral pH (probably through transcription of target genes). Thus, *SNF5* appears to both contribute to pH regulation and respond to pH changes through its polyQ domain and histidines therein.

### *ADH2* induction requires transient intracellular acidification

The dynamics of pH_i_ and *P_ADH2_-mCherry* suggested that intracellular acidification at the beginning of carbon starvation might be required to induce *ADH2* expression. To test this hypothesis, WT cells were subjected to glucose-starvation for 2 hours at pH_env_ 5 and then the pH_env_ was alkalinized to 7.4, thus forcing neutralization of the cytosol (supplemental figure 6A). pHluorin measurements confirmed that pH_i_ of the entire population was forced to 7 by this treatment (supplemental figure 6C). The transient acidification during the first 2 hours of starvation was sufficient to allow robust induction of *P_ADH2_-mCherry* (supplemental figure 6B, D). The reverse experiment, where cells are transiently glucose-starved at pH_env_ 7.4 for 2 h and then switched to pH_env_ 5 did not lead to efficient induction of *P_ADH2_-mCherry*, despite the pH_i_ drop when switched to pH_env_ 5 after the 2^nd^ hour (supplemental figure 6B, D). We also used sorbic acid, a weak organic acid that can shuttle protons across the plasma membrane (Munder *et al*., 2016) to prevent pH_i_ recovery during glucose starvation. As expected the entire population remained acidic through the course of the experiment and no *P_ADH2_-mCherry* induction was observed (supplemental figure 7) confirming that recovery of neutral pH_i_ is also required for *ADH2* induction. Together, these results indicate that *ADH2* induction requires a transient drop in pH_i_ at the beginning of glucose-starvation treatment followed by recovery to neutral pH_i_.

### Transient acidification also occurs in response to oxidative stress

We posited that the polyQ domain of *SNF5* could sense pH_i_ changes as a more general mechanism for transcriptional reprograming upon environmental changes. After screening several more stresses, we observed that *ΔQ-snf5* had very poor fitness in media supplemented with 1 mM peroxide (supplemental figure 8A). Motivated by this result, we tested if oxidative stress induce changes in pH_i_. Indeed, similarly to carbon starvation, addition of peroxide to the media led to acidification of the cytosol to pH ~ 6.5 (supplemental figure 9B, first panel). Also like carbon starvation, this pH_i_ drop didn’t occur in the whole population: About 40% of the population acidified after a 30 min incubation with 1 mM peroxide. These cells recovered to neutral pH with similar kinetics to cells that eventually induce *ADH2* following carbon starvation. The degree of acidification upon peroxide treatment is dose-dependent with an EC50 around 1 mM – at which point the pH response of the population is bimodal, also similar to carbon starvation. *SNF5* mutants had markedly different pH responses. The *snf5Δ* strain failed to acidify its cytoplasm, while the *ΔQ-snf5* and *_HtoA-_snf5*, alleles acidified normally but then failed to recover neutral cytosolic pH (supplemental figure 8B). This striking similarity of the cytosolic pH responses in acute starvation, recovery from starvation, and oxidative stress suggests a general pH driven mechanism for transcriptional reprogramming. The equivalent effects of the WT, snf5Δ, *ΔQ-snf5* and *_HtoA-_snf5* alleles during all of these state changes highlights the importance of the *SNF5* polyQ domain in pH regulation.

### *SNF5* polyQ and an acidic environmental pH are required for the efficient induction of a large number of glucose-repressed genes

We performed RNA sequencing analysis on total mRNA extracted from WT, *snf5Δ, ΔQ-snf5* and *_HtoA-_snf5* strains during exponential growth (+Glu) and after 4 hours of glucose starvation at pH_env_ 4 (optimal starvation condition). To determine the effect of environmental pH, we also analyzed mRNA content of WT strains starved for glucose at pH_env_ 7. Principal component (PC) analysis was performed to extract the major differences between conditions. Glycolysis and glucose metabolism, and peroxisome and carnitine metabolism genes dominate the PC1 axis. Similarly, PC2 contains genes involved in sugar import at one end, and microbody and fatty acid degradation at the other, consistent with a shift from carbon rich anabolic processes to catabolism during starvation (figure 4A). All biological replicates clustered well on the y-axis (PC2), and most of them did so on the x-axis (PC1), confirming the reproducibility of our results. However, replicates of WT, *ΔQ-snf5* and *_HtoA-_snf5* strains showed variability on the PC1 axis during glucose starvation. We hypothesize that bistability in the expression of glucose-repressed genes and high sensitivity to minor environmental or epigenetic variations could be responsible for this greater variability on PC2.

**Figure 4:**
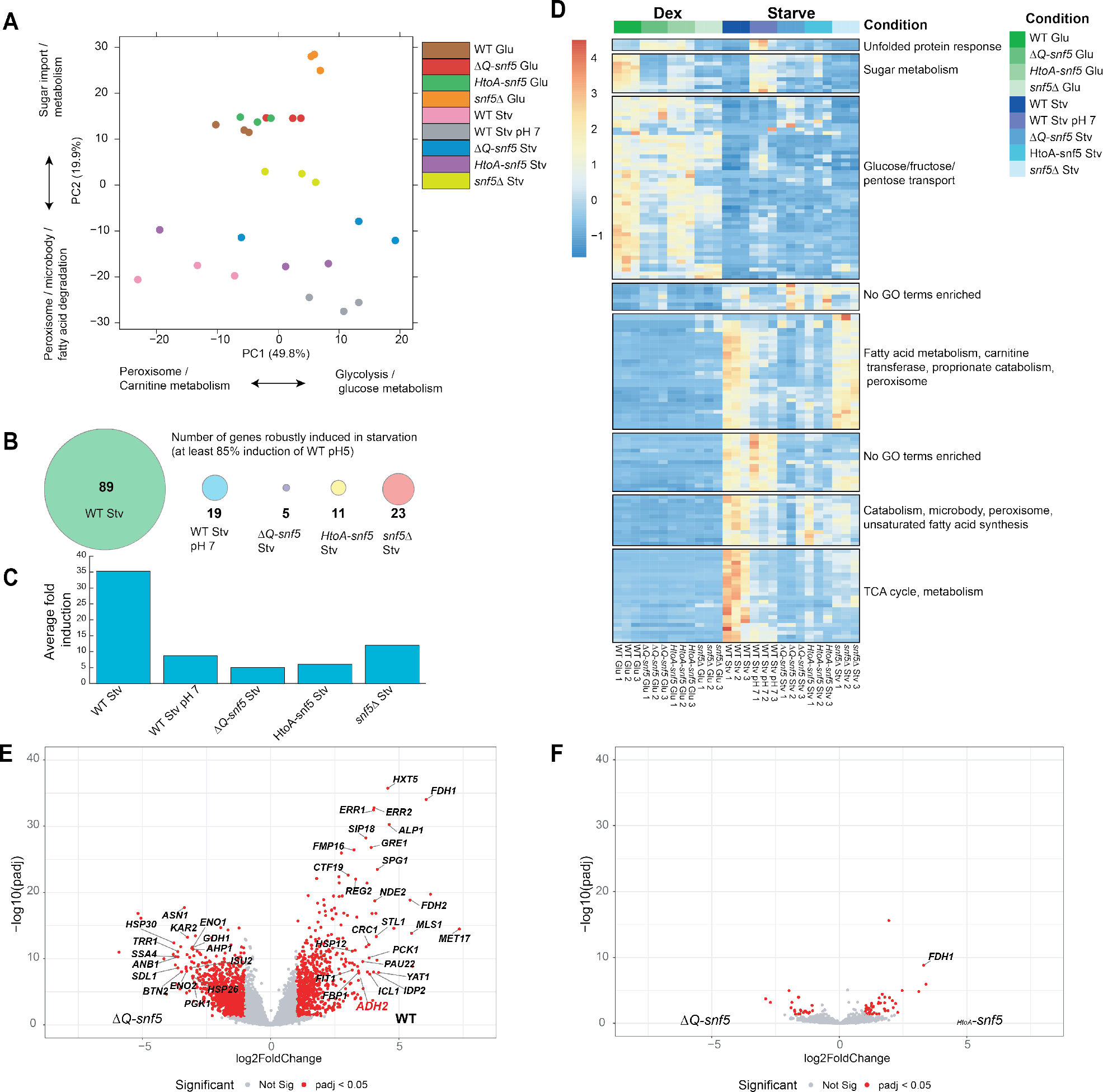
RNA-seq analysis reveals widespread polyQ-dependence in the expression of glucose repressed genes. A- Principal component (PC) analysis of 3 replicates in each condition tested. B- Number of glucose repressed genes induced to at least ~85% of WT induction in each condition. C- Average Fold induction of glucose-repressed genes upon glucose-starvation. D- Heat map of row and column normalized expression values of 149 genes significantly differently expressed between WT Glu and WT Starve (log2 fold change = 1 and adjusted p value = 0.05 (Wald test)). (Right) Gene ontology enrichment results for 8 clusters of genes with similar expression signatures across conditions. E and F- Volcano plots of genes with significantly different expression in starvation conditions between *ΔQ-snf5* and WT (left) and *_HtoA-_snf5* (right) (log2 fold change = 1 and adjusted p value = 0.05 (Wald test)

We defined a set of 149 genes that were differentially expressed (> 2 fold change at an adjusted p-value of < 0.05, Wald test) in wild type cells in logarithmic growth in glucose compared to 3 h of carbon starvation at pH_env_: 4. Around half of these genes (89 genes) were induced during starvation. Of this set of glucose-repressed genes, only 19 (21%) retained expression (at > 85% WT level) when the WT strain was starved at pH_env_ 7, and only 5 (6%) were induced in the *ΔQ-snf5* (figure 4B). WT strains upregulated these genes an average of 35-fold in optimal conditions, while gene induction was < 10-fold in starvation of WT cells at pH_env_ 7.5 and in *SNF5* mutant alleles (figure 4C).

Hierarchical clustering of the differentially expressed genes (Euclidean distance) revealed some differences in gene expression between *SNF5* alleles in logarithmically growing cultures. For example, the mRNAs of genes related to sugar metabolism and transport generally decreased in *SNF5* mutant alleles (figure 4D, top). However, there was far greater variability in carbon starvation. Gene ontology (GO) analysis for genes that failed in all *SNF5* mutants and WT strains in starvation at pH_env_ 7, showed enrichment of Krebs cycle, carbon metabolism and glycolysis/gluconeogenesis genes (figure 4D). Genes involved in fatty acid metabolism and carnitine transport failed to induce in WT strains starved at pH_env_: 7, *ΔQ-snf5*, and *_HtoA-_snf5* strains (figure 4D, middle). However, expression of these genes still occurs in *snf5Δ* strains (figure 4D, bottom). This may explain why *SNF5* has not previously been linked to these processes.

Volcano plot analysis show marked global differences in transcription of WT and *ΔQ-snf5* strains during glucose starvation (figure 4E), while *ΔQ-snf5* and *_HtoA-_snf5* are very similar (figure 4F), confirming that the polyQ of *SNF5* plays an important regulatory role for a large number of genes and that the histidines within this domain are crucial determinants of this regulation. Together, these data suggest that our previous observations using *ADH2* as a reporter of glucose-repressed genes can be generalized more broadly to the carbon starvation response.

### Spidroin partially complements the *SNF5* polyQ domain

Given the ability of polyQs to form aggregates, we hypothesized that pH may induce changes in the structural or oligomerization state of Snf5 and therefore the SWI/SNF complex. Genetic analysis of GFP tagged *SNF5* and the mutant strains led us to the conclusion that GFP interferes with proper folding of the SWI/SNF complex (data not shown). Therefore, unable to directly visualize *SNF5* by fluorescence microscopy, we tried an orthogonal way to test our hypothesis. To this end, we took advantage of the spidroin protein. Spidroin, the main component of spider silk, is made and stored in the silk glands of spiders in soluble form at pH ≥7. The pH of the silk duct decreases from around 7 at the proximal end to ≤6.3 at the distal end causing spidroin to oligomerize into a silk fiber (Askarieh *et al*., 2011) (figure 5A). Thus, we engineered a synthetic minispidroin allele of 376 amino acids in length (similar to the size of the polyQ domain) to use as an alternative pH-sensing domain.

**Figure 5:**
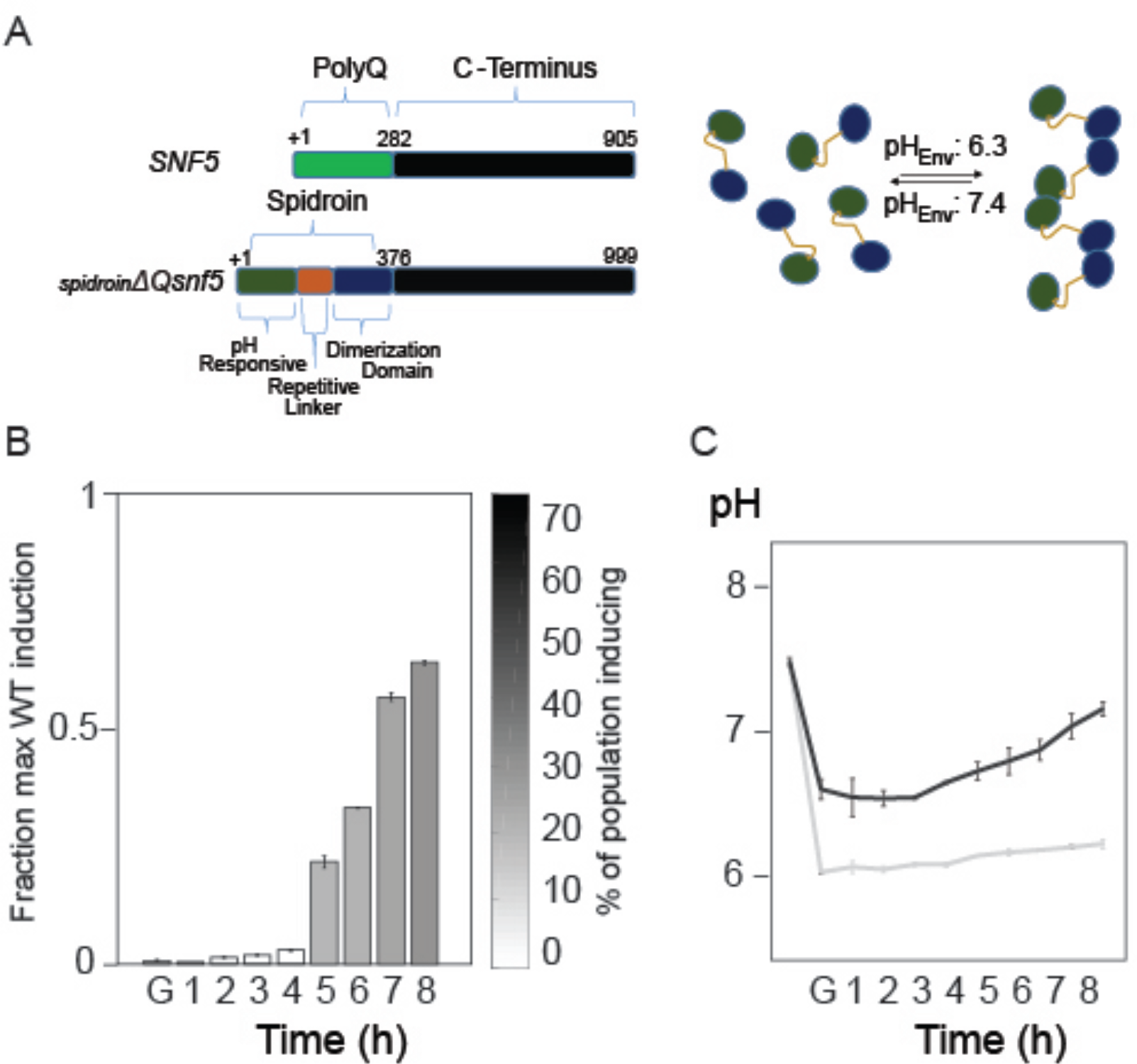
A synthetic spidroin domain partially rescues pH-dependent induction of transcription. A- Schematic representation of *Spidroin-ΔQ-snf5* and its aggregation behavior. B- Quantification of *P_ADH2_-mCherry* expression on *_Spidroin_-ΔQ-snf5* strain at various time points after glucose withdrawal. The y-axis is the intensity of fluorescent reporter (normalized to the levels of wild type control cells), shading corresponds to the percentage of cells inducing at each time point. D- Quantification of cytosolic pH.

First, we tested if acidification of the cytosol upon glucose starvation could induce aggregation of the spidroin peptide. Exponentially growing cells with a cytosolic pH of 7.4 did not show any spidroin aggregation. We then manipulated the pH_i_ by poisoning glycolysis and the electron transport chain with 2-deoxyglucose (2-DG) and sodium azide (AZ) respectively. This treatment leads to depletion of ATP causing proton pumps to fail, thereby leading to equilibration of the cytosolic pH with the pH_env_. Spidroin aggregated in over 90% of cells when the pH_i_ was reduced to 5.5 (supplemental figure 10A, B). However, no aggregation was seen in cells treated with 2-DG and AZ at pH_env_ 7.5 (supplemental figure 9A, B). These results indicate that our synthetic mini-spidroin domain changes aggregation state in a pH-dependent manner when expressed in *S. cerevisiae*. This aggregation also occurred during glucose starvation: At pH_env_ 5.5 around 40% of the cells have clear aggregates, while at pH_env_ 7.5 only about 5% do (supplemental figure 9B, C). These results are consistent with our pH_i_ measurements during glucose-starvation.

After confirming that at least 40% of the culture would induce spidroin aggregation upon glucose-starvation at pH_env_ 5.5, we fused the mini-spidroin domain to the N-terminus of the *ΔQ-snf5* gene, thus replacing the polyQ of *SNF5* with the spidroin peptide (figure 5A). Amazingly, *_Spidroin_-ΔQ-snf5* partially rescued the defects observed in the *ΔQ-snf5* strain (figure 5B, C, and supplemental figure 3). After a transient acidification ~ 50% of the population induced *ADH2* with mean induction levels reaching 65% of WT control levels (figure 5B). As expected *ADH2* expression was pH sensitive in the *_Spidroin_-ΔQ-snf5* strain with a pH_env_ optimum of 5.5, and no induction when pH_env_ was buffered to 6.5 or above (figure 5B, supplemental figure 3). Thus, *_Spidroin_-ΔQ-snf5* partially recapitulates the *ADH2* expression and the cytosolic pH dynamics of WT.

We evaluated whether other sequences known to undergo phase transitions, but not affected by pH, could complement *SNF5* polyQ function. We fused the low complexity domains of the FUS and hnRNP proteins to the N-terminus of *ΔQ-snf5*. These domains have been reported to form hydrogels that can recruit RNA Polymerase II (RNA-polII) (Kato *et al*., 2012; Kwon *et al*., 2013). However, these domains did not complement polyQ function: *_FUS_-ΔQ-snf5* and *_hnRNP_-ΔQ-snf5* both completely failed to induce *P_ADH2_*-mCherry expression (data not shown). This result suggests that a pH-responsive domain is required for SWI/SNF function.

## Discussion

Dynamic changes in cytosolic pH have been observed in several organisms. pH regulation occurs in many physiological contexts including: cell cycle progression (Gagliardi and Shain, 2013); the circadian rhythm of crassulacean acid metabolism plants (Hafke *et al*., 2001); oxidative stress (van Schalkwyk *et al*., 2013); temperature or osmotic shock (Karagiannis and Young, 2001); and changes in nutritional state in *Saccharomyces cerevisiae* (Orij *et al*., 2009). However, the physiological role of these pH_i_ fluctuations remains poorly understood. Our work establishes a functional link between cytosolic pH changes and gene expression.

Previous studies reported population averages of pH_i_ (Orij *et al*., 2009) and have emphasized the inactivation of processes in response to cytosolic acidification (Petrovska *et al*., 2014; Joyner *et al*., 2016; Munder *et al*., 2016). However, it is unclear how necessary modifications to the cell can occur if cellular dynamics are uniformly decreased. Much less has been reported regarding a potential positive role of fluctuations in pH_i_. We found that transient acidification is indeed required for activation of a set of stress response genes.

Using single cell measurements of both pH_i_ and gene expression we found that two subpopulations arose upon acute glucose-starvation, one with pH_i_ ~5.8 and a second at ~6.2. The latter population recovered neutral pH_i_ and then induced glucose-repressed genes, while the former population remained dormant. We have not yet determined the mechanism that drives the bifurcation in pH response but we did not find any correlation to cell cycle stage or cell age (data not shown). We speculate that bistability may provide a form of bet hedging (Levy, Ziv and Siegal, 2012) where some cells attempt to respond to carbon starvation, while others enter a dormant state (Munder *et al*., 2016).

It is becoming clear that pH is an important mechanism of biological control. It was previously shown that the protonation state of phosphatidic acid (PA) determines binding to the transcription factor Opi1, coupling membrane biogenesis and intracellular pH (Young *et al*., 2010). We focused our studies on the N-terminal region of *SNF5* because it is known to be important for the response to carbon starvation and contains a large low complexity region enriched in both glutamine and histidine residues. Histidines are good candidates for pH sensors as they can change protonation state over the recorded range of physiological pH. Indeed, the polyQ domain and the histidines contained within were required for both transcriptional reprogramming and for regulation of intracellular pH. It also seems that the *SNF5* polyQ plays a more general role as a pH sensor as we observed defects in pH_i_ regulation in the *ΔQ-snf5* and *snf5Δ* in response to oxidative stress.

PolyQs have been identified as the molecular basis of nine neurodegenerative diseases (Fan *et al*., 2014) and have been mostly studied as seeds of toxic aggregation. The aggregation state of polyQs has been linked to the length of the homorepeat (Bates *et al*., 2015; Kuiper *et al*., 2017). However, recently published evidence indicates that the nature of the residues at the boundaries of polyQs can also strongly impact aggregation propensity. For instance, a proline homorepeat at the C-terminal of the Huntingtin protein-polyQ prevents aggregation while 17 residues with an overall positive charge at the N-terminus have the opposite effect, inducing stronger aggregation at neutral pH (Thakur *et al*., 2015; Shen *et al*., 2016). Similarly, the addition of two negatively charged residues at the amino end and two positively charged residues at the carboxyl end of the polyQ domain of the yeast Sup35 solubilizes the protein at acidic pH and aggregates at neutral pH while a polyalanine flanked by the same charged amino acid is soluble at all pH values (Perutz *et al*., 2002). We hypothesize that the protonation state of histidines within or at the boundaries of polyQs regulates their aggregation state. The precise nature of this state remains unclear but could range from an oligomerization to a phase transition similar to those that have been recently described for many other intrinsically disordered domains and proteins (Brangwynne, Mitchison and Hyman, 2011; Kato *et al*., 2012; Kwon *et al*., 2013) including polyQ proteins (Zhang *et al*., 2015). We could not detect large phase separated structures by fluorescence microscopy of Snf5p during glucose-starvation (data not shown). However, it has been recently shown that polyQs made of 34 to 96 glutamines convert between three states: soluble protein, small rapidly-diffusing clusters and large slowly-diffusing aggregates (Li *et al*., 2016). It is possible that the SWI/SNF complex converts to clusters that are below the diffraction limit of conventional microscopy. Single molecule microscopy (Li *et al*., 2016) or other super-resolution techniques will be required to directly observe these events, which are likely to be transient in nature and exist only briefly. Nevertheless, we were able to design experiments to address this hypothesis using the spider silk protein, spidroin.

Spidroin is a spider web peptide that is soluble at pH 7 and aggregates at pH 6.3 (Askarieh *et al*., 2011). Importantly, spidroin does not contain polyQ repeats. Thus, spidroin is an orthogonal pH-dependent domain that dynamically changes aggregation states over a similar range of conditions to those sampled during carbon starvation in yeast. This domain allowed us to test our hypothesis that the *SNF5* polyQ undergoes pH-dependent aggregation: If the *ΔQ-snf5* allele is defective due to a failure to aggregate, then adding the synthetic spidroin domain should revert this phenotype. Indeed, we observed that *ADH2* expression was partially recovered in the *_Spidroin_-ΔQ-snf5* strain. Therefore, we hypothesize that transient aggregation is part of the mechanism by which the SWI/SNF complex drives transcriptional reprogramming.

The “aggregated” polyQ could actually be more than one state – for example, solid gel-like and viscous liquid-like behaviors could exist. The fact that acidification followed by neutralization is required for efficient transcriptional reprogramming suggests that the physical changes in SWI/SNF are transient, but there must be some memory of this event. Several models could account for this memory. For example, transient physical state changes could serve to redistribute the SWI/SNF complex to new loci, change the composition of “super-complexes” of SWI/SNF with other molecules in the nucleus, or recruit enzymes to changes the post-translational modifications on SWI/SNF. Further studies will be required to elucidate the details of the transcriptional reprogramming mechanism.

During glucose starvation, many SWI/SNF-dependent genes are inactivated and a new set of genes is induced. We speculate that the *SNF5* polyQ domain acts as a “reset” switch to allow this redistribution of the SWI/SNF complex upon starvation. The pH dependent genes that require the SNF5 polyQ are enriched in fatty acid metabolism. This could be particularly important in cancer where it has been observed that about 20% of human cancers have mutations in the SWI/SNF complex (Kadoch *et al*., 2013). Human *SNF5* (SMARCB1) was the first subunit of the SWI/SNF to be linked to cancer, where it is mutated in most cases of pediatric malignant rhabdoid tumor (Biegel *et al*., 1999; Sevenet *et al*., 1999). It is known that mutations of the SWI/SNF that lead to cancer generally result in missregulation of fatty acid synthesis, which is required for cancer proliferation (Wu *et al*., 2016; Nickerson *et al*., 2017). However, how pH and human homologs of yeast SWI/SNF polyQs affect expression of these genes has yet to be investigated. An acidic pH is a prominent feature of the tumor microenvironment (Wike-Hooley, J. Haveman and Reinhold, 1984; Tannock and Rotin, 1989), raising the possibility that SNF5 may play a pH-sensing role in human disease.

All cells must modify gene expression to respond to environmental changes. This phenotypic plasticity is essential to all life, from single celled organisms fighting to thrive in a competitive environment, to the complex genomic reprogramming that must occur during development and tissue homeostasis in the *metazoa*. Despite the differences between these organisms, the mechanisms that regulate gene expression are highly conserved. This work provides new insights into pH as a signal through which life perceives and reacts to its environment.

**Table 2.** Strains used in this study.

|  |
| --- |
| *ura3Δ0 his3Δ0 leu22Δ0 met15Δ0 SNF5* |
| *ura3Δ0 his3Δ0 leu22Δ0 met15Δ0 ΔQ-SNF5::KAN* |
| *ura3Δ0 leu22Δ0 met15Δ0 HtoA-SNF5::HIS3* |
| *ura3Δ0 leu22Δ0 met15Δ0 SpidroinSNF5::HIS3* |
| *his3Δ0 leu22Δ0 met15Δ0 SNF5 pADH2-mCherry::URA3* |
| *his3Δ0 leu22Δ0 met15Δ0 ΔQ-SNF5::KAN pADH2-mCherry::URA3* |
| *leu22Δ0 met15Δ0 HtoA-SNF5::HIS3 pADH2-mCherry::URA3* |
| *leu22Δ0 met15Δ0 SpidroinSNF5::HIS3 pADH2-mCherry::URA3* |
| *his3Δ0 met15Δ0 SNF5 pADH2-mCherry::URA3 pHluorin::LEU2* |
| *his3Δ0 met15Δ0 ΔQ-SNF5::KAN pADH2-mCherry::URA3 pHluorin::LEU2* |
| *met15Δ0 HtoA-SNF5::HIS3 pADH2-mCherry::URA3 pHluorin::LEU2* |
| *met15Δ0 SpidroinSNF5::HIS3 pADH2-mCherry::URA3 pHluorin::LEU2* |
| *ura3Δ0 his3Δ0 LEU22Δ0 met15Δ0 SNF5::KAN (CEN/ARS-SNF5::URA3)* |
| *ura3Δ0 his3Δ0 met15Δ0 snf5Δ::KAN (CEN/ARS-SNF5::URA3) pADH2-mCherry::LEU2* |
| *ura3Δ0 his3Δ0 met15Δ0 snf5Δ::KAN (CEN/ARS-SNF5::URA3) pADH2-mCherry::LEU2 pHluorin::NAT* |
| *leu22Δ0 met15Δ0 pADH2-mCherry ::URA3 SNF2-TAP::HIS3* |
| *leu22Δ0 met15Δ0 ΔQ-SNF5::HIS3 pADH2-mCherry::URA3 SNF2-TAP::KAN* |
| *leu22Δ0 met15Δ0 HtoA-SNF5::HIS3 pADH2-mCherry::URA3 SNF2-TAP::KAN* |
| *his3Δ0 met15Δ0 (CEN/ARS-SNF5::URA) SNF5::KAN pADH2-mCherry::LEU2 SNF2-TAP::NATMX* |
| *his3Δ0 met15Δ0 SNF5::KAN pADH2-mCherry::LEU2 (CEN/ARS-SNF5::URA3)* |
| *his3Δ0 met15Δ0 ΔQ-SNF5::KAN pADH2-mCherry::LEU2 (CEN/ARS-SNF5::URA3)* |

## Supplemental figures

**Supplemental figure 1:**
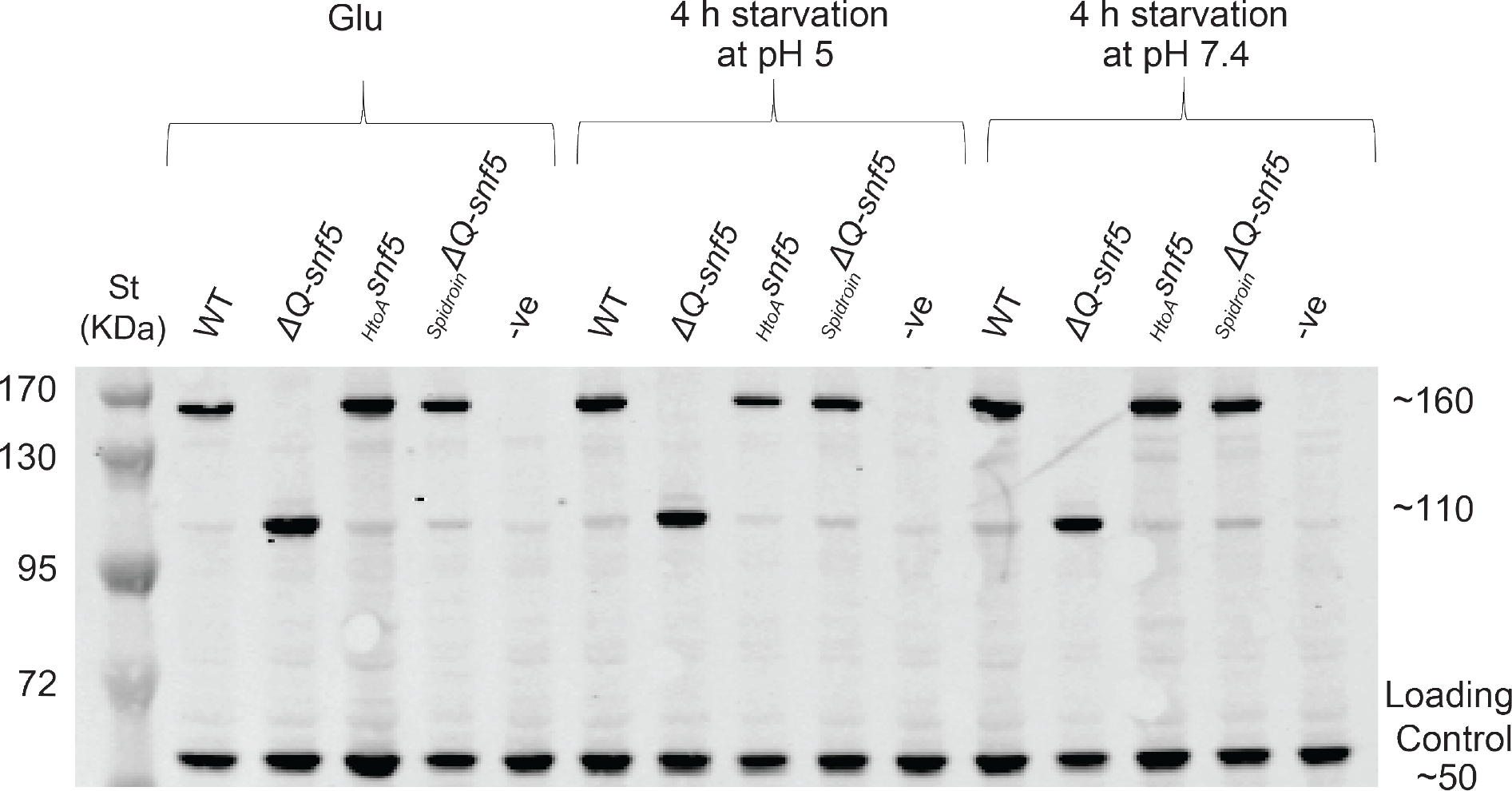
Neither *SNF5* nor its mutant alleles are degraded upon glucose-starvation. Western blots of the *SNF5-TAP* and TAP tagged mutant alleles. *ΔQ-snf5* is 288 amino acids smaller than WT. Anti-glucokinase antibody was used as a loading control (Bottom band at ~50 kDa).

**Supplemental figure 2:**
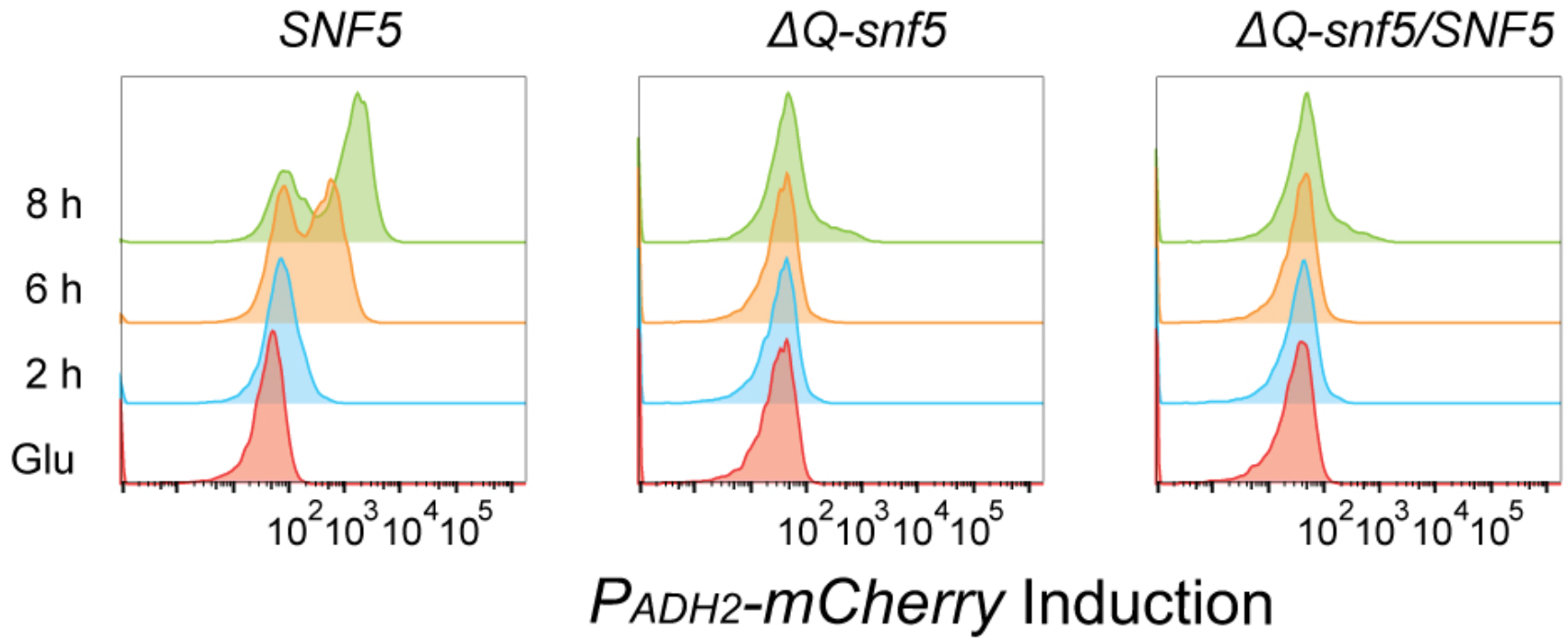
Deletion of *SNF5* polyQ creates a dominant negative allele. Histograms of *P_ADH2_-mCherry* reporter expression, y-axis is time in hours, starting immediately prior to glucose-starvation (“Glu”), WT/*ΔQ-snf5* indicates a strain carrying both alleles integrated at the endogenous locus

**Supplemental figure 3:**
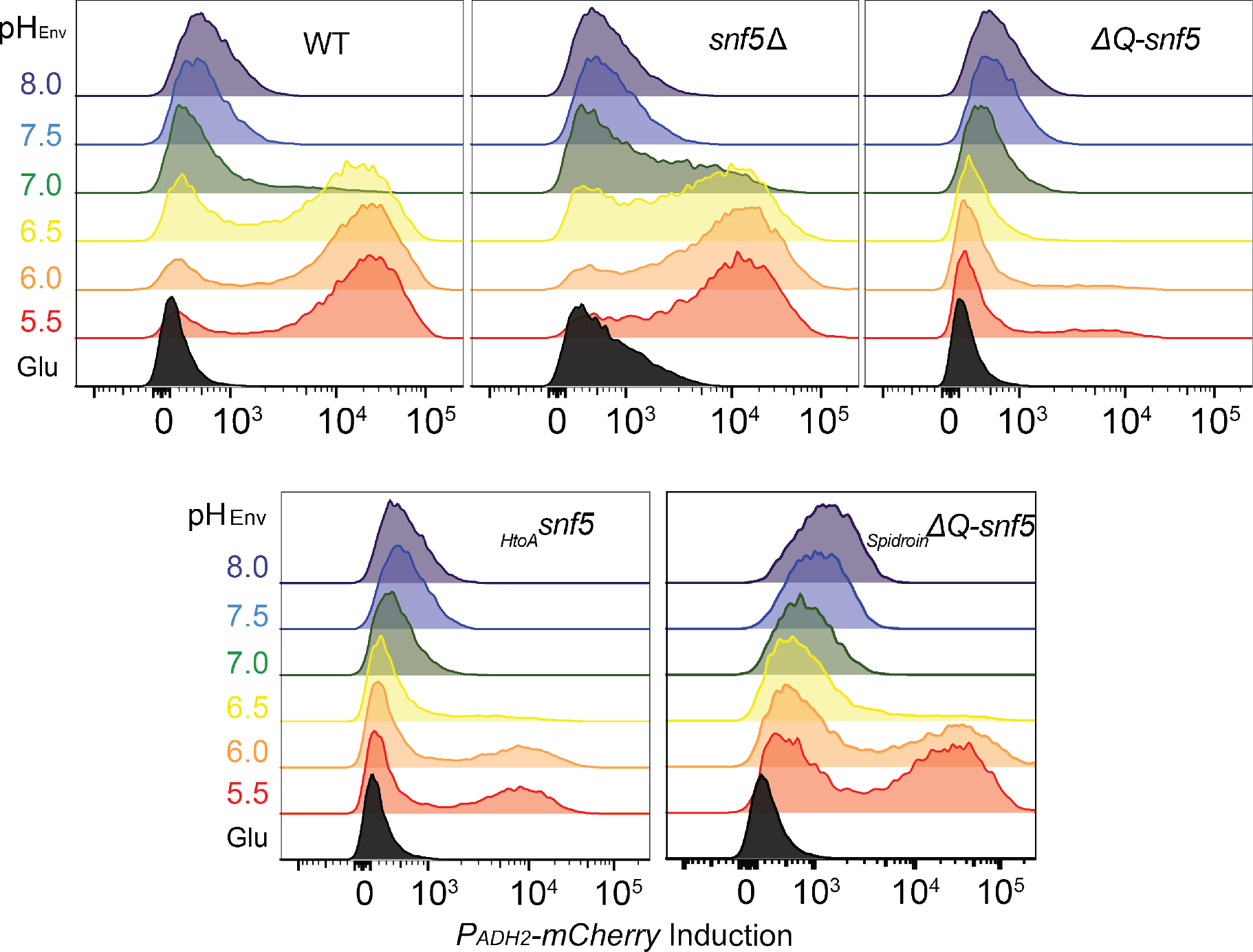
*ADH2* expression is pH-dependent. Cytometry data histograms of *P_ADH2_-mCherry* induction. Environmental pH is indicated by color codes: pH 5.5 (red); 6.0 (orange); 6.5 (yellow); 7.0 (green); 7.5 (blue); and 8.0 (purple).

**Supplemental figure 4:**
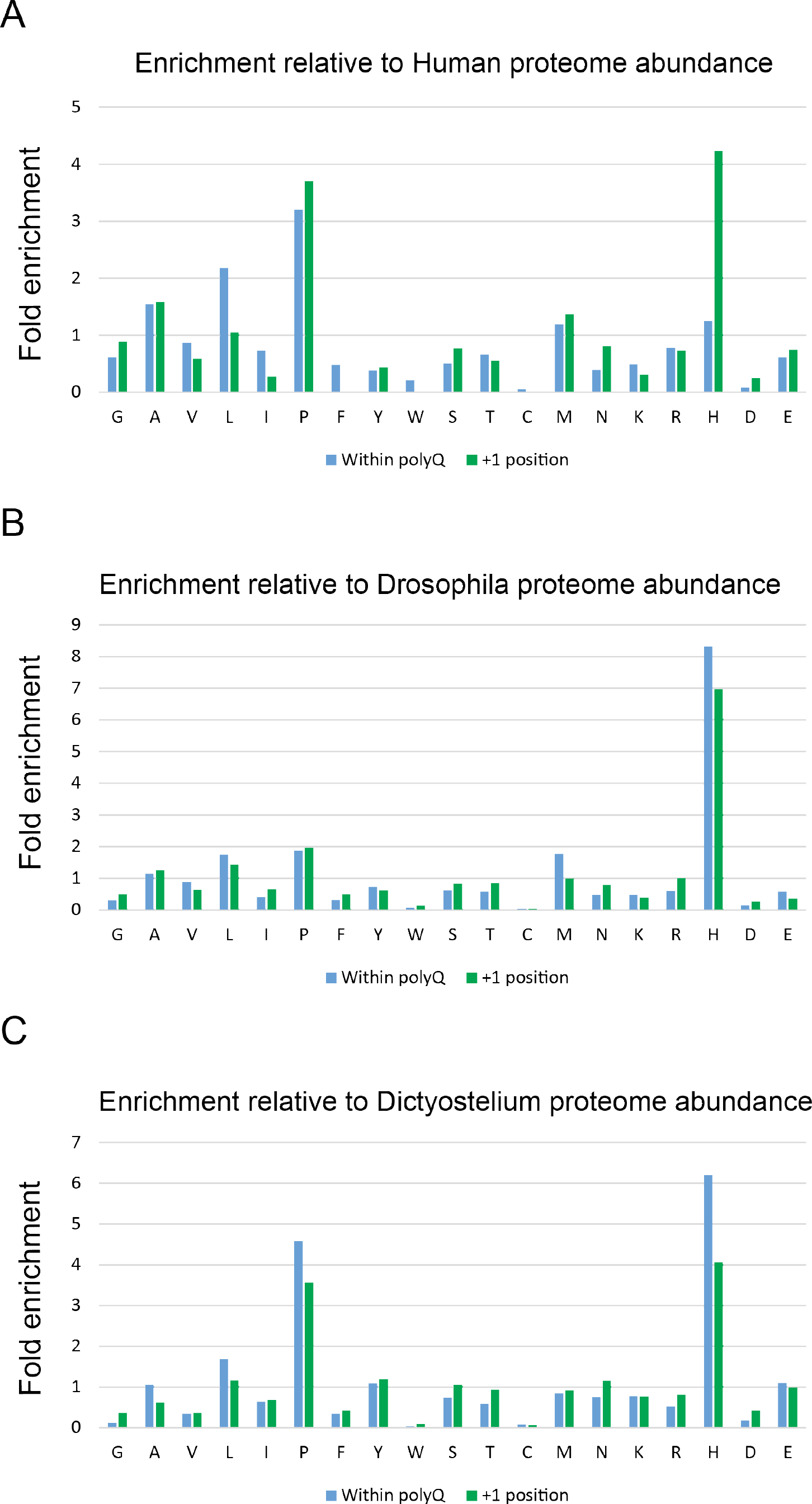
A- Sequence analysis results for all Human polyQs, showing frequencies of each amino acid within polyQ structures (blue), and immediately C-terminal to the structures (the +1 position, green). B- Same analysis for all Drosophila polyQs. C- Same analysis for all Dictyoselium polyQs.

**Supplemental figure 5:**
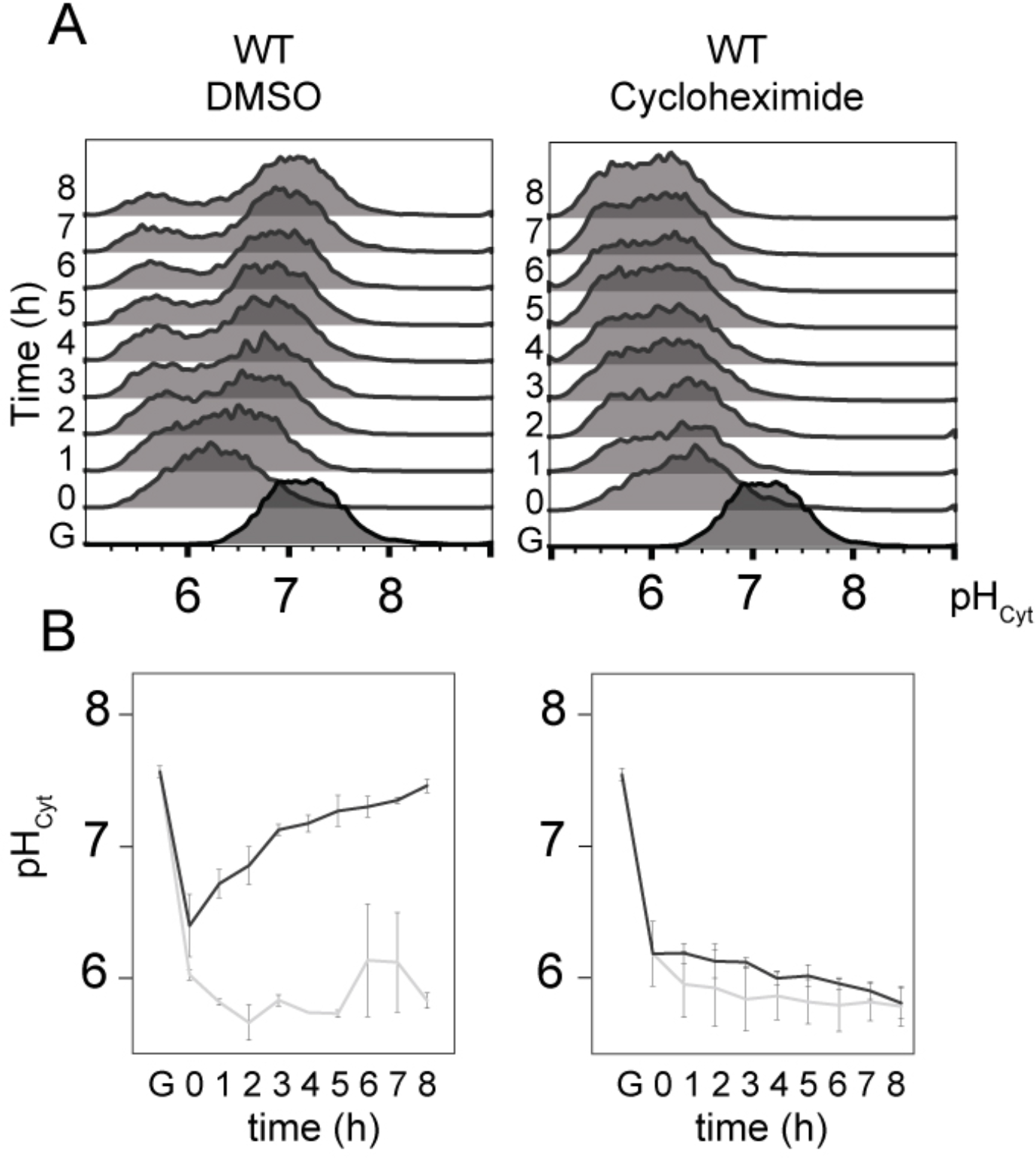
pH_i_ recovery during glucose-starvation requires translation. A- Histograms of cytosolic pH upon glucose-starvation of WT strains treated with 35 mM Cycloheximide and control. B- Quantification of the data plotted in A.

**Supplemental figure 6:**
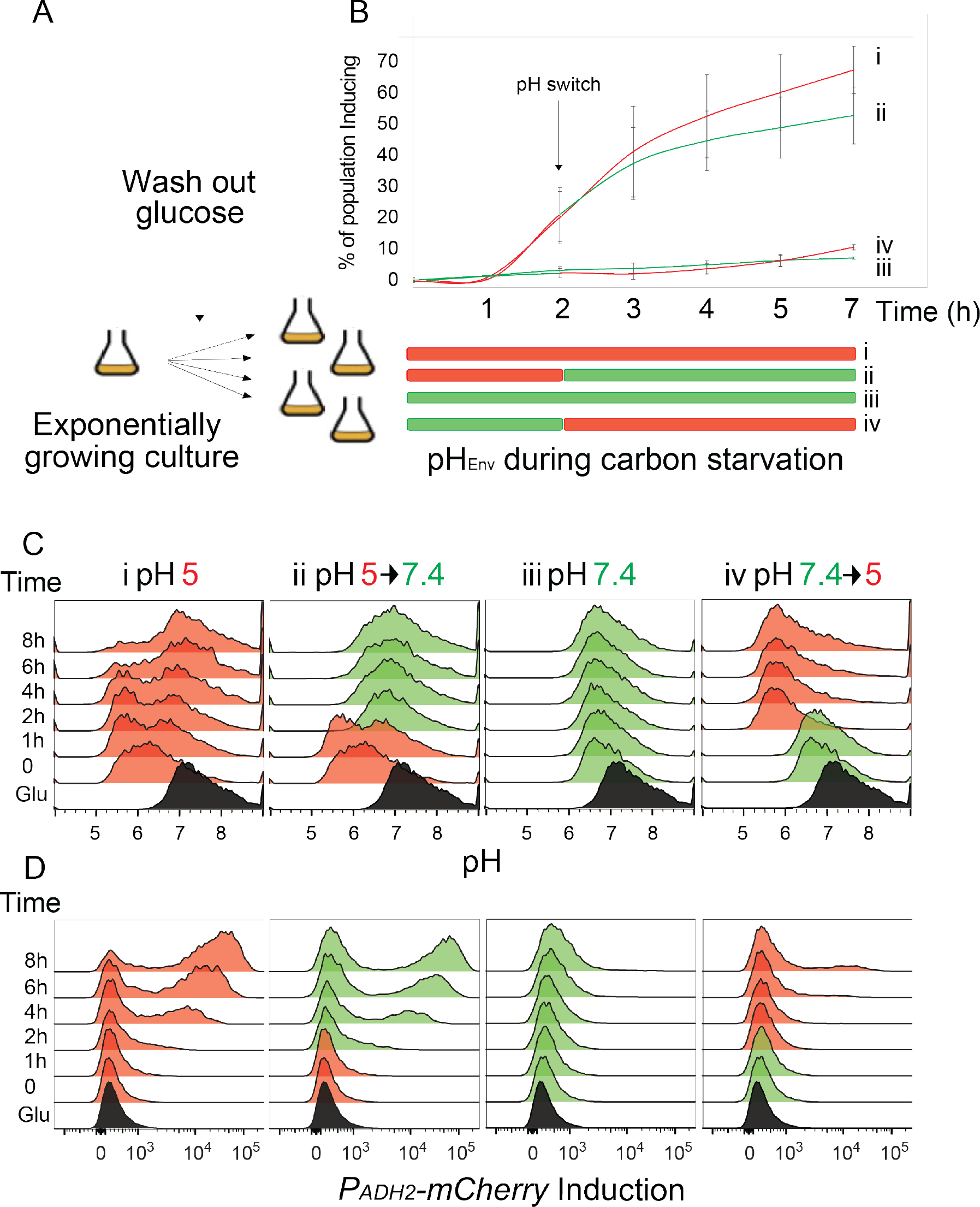
Transient acidification of the cytosol is sufficient for *ADH2* expression. A- Outline of the experiment, exponentially growing cultures are divided in four: two of them washed into glucose-starvation media at pH: 5 (red), and the other two into starvation media at pH: 7.4 (green). After two hours of glucose-starvation, one culture at pH_env_ 5 was switched to 7.4 and one culture that was at pH_env_ 7.4 was switched to 5. B- *P_ADH2_-mCherry* reporter expression, x-axis is time in hours (pH switch occurred at 2 h), y-axis is the percentage of cells that induced the *P_ADH2_-mCherry* reporter. C- Cytosolic pH histograms, culture in complete media is labeled “Glu”, red histograms indicate cultures starved at pH_env_ 5, and green histograms indicate cultures starved at pH_env_ 7.4, y-axis is time in hours. D- Histograms of *P_ADH2_-mCherry* reporter expression over time, color-coded as in C.

**Supplemental figure 7:**
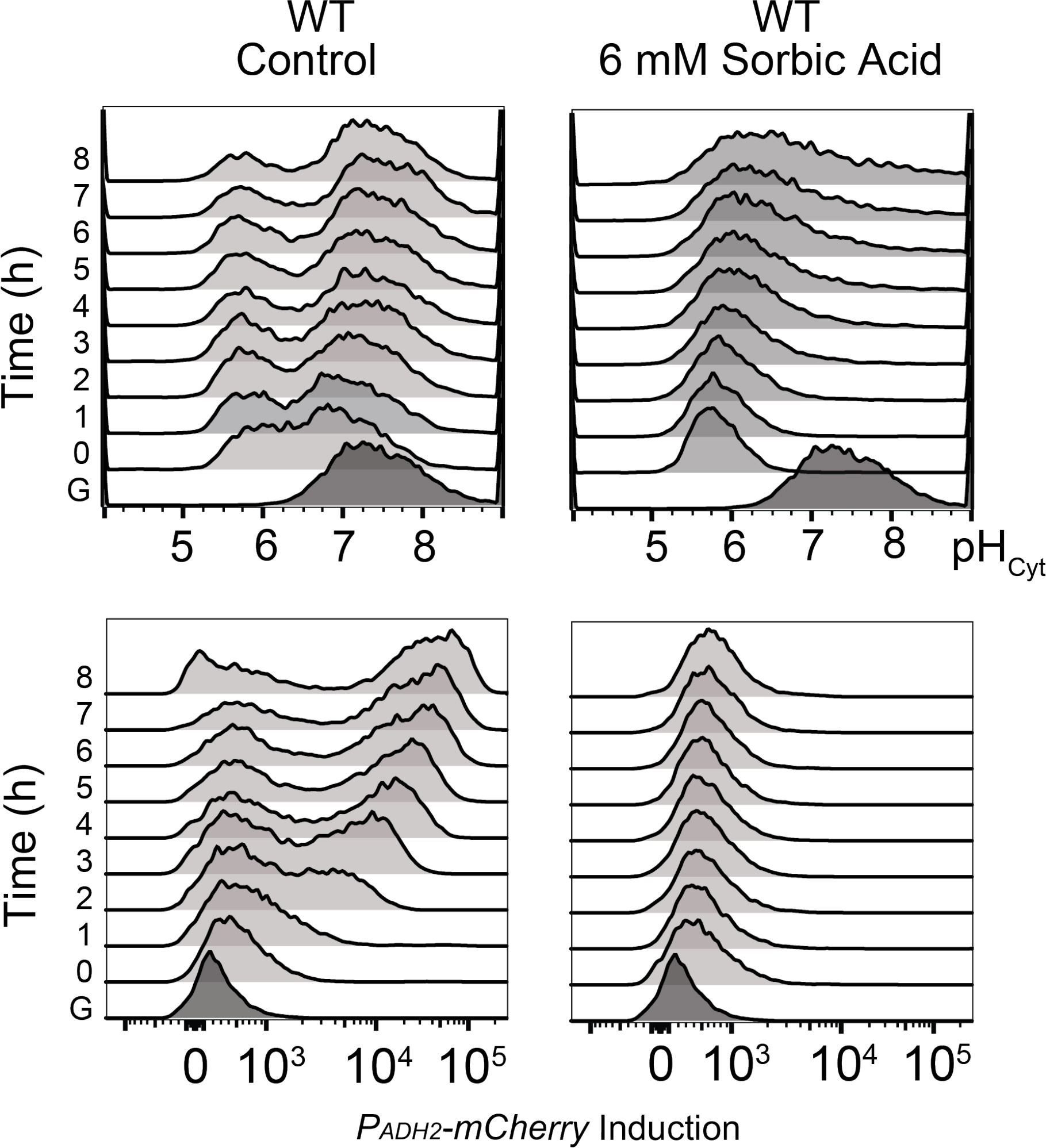
pH_i_ recovery is required for efficient induction of *ADH2* reporter. Histograms of cytosolic pH and *P_ADH2_-mCherry* induction upon glucose starvation on control media and supplemented with 6 mM sorbic acid.

**Supplemental figure 8:**
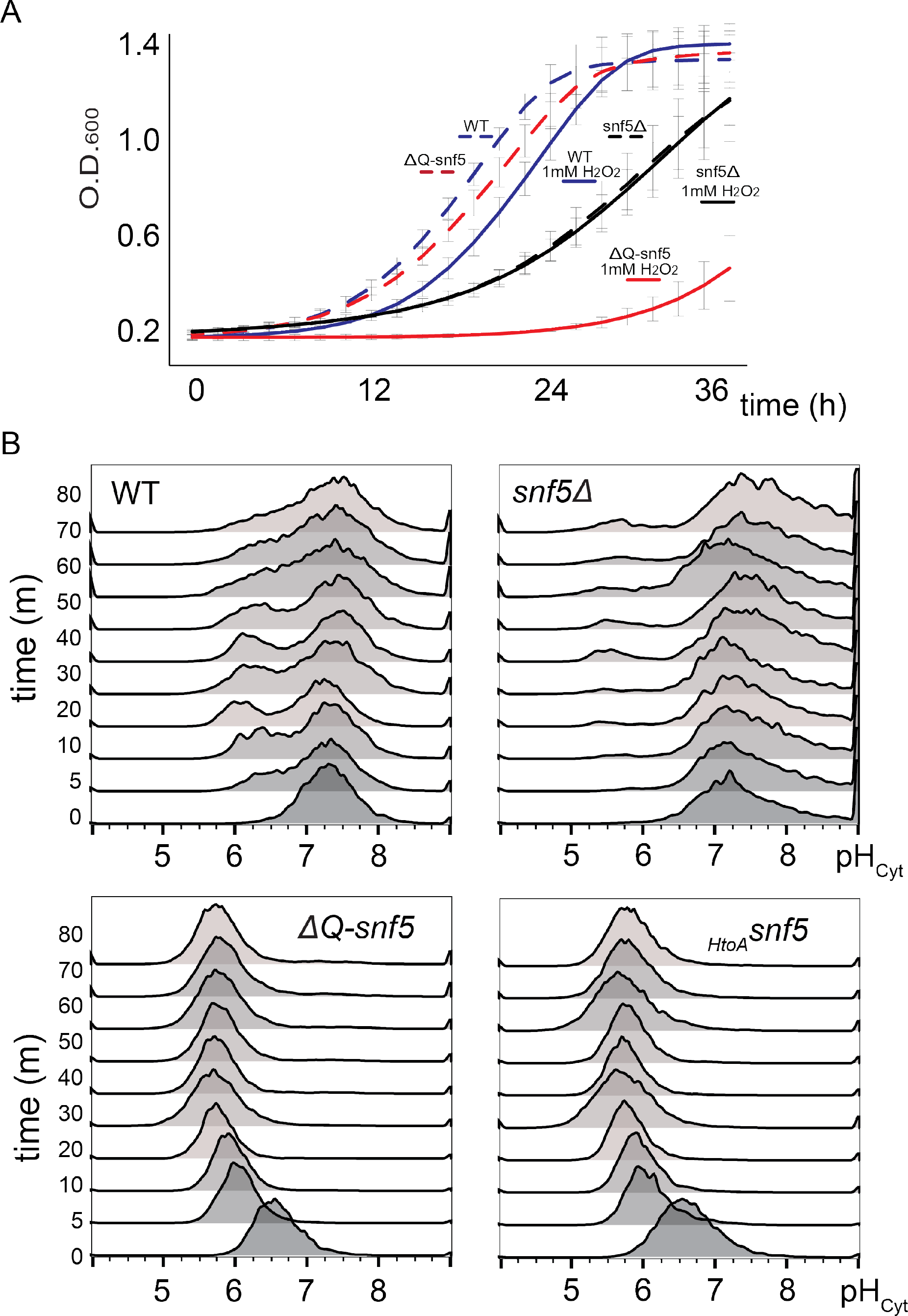
Phenotypes of *SNF5* mutants under oxidative stress. A- Growth curves of WT, *ΔQ-snf5* and *snf5Δ* strains. From left to right, WT (dashed blue), *ΔQ-snf5* (dashed red), WT 1mM H2O2 (blue) *snf5Δ* (dashed black) *snf5Δ* 1 mM H_2_O_2_ (black), *ΔQ-snf5* 1 mM H_2_O_2_ (red). Exponentially growing cultures OD_600_: ~0.3 were inoculated (final OD_600_: 0.01) with and without 1 mM H_2_O_2_. B- Histograms of cytosolic pH of WT, *snf5Δ, ΔQ-snf5* and *_HtoA-_snf5* upon addition of 1 mM H_2_O_2_ from Time 0 to 80 min.

**Supplemental figure 9:**
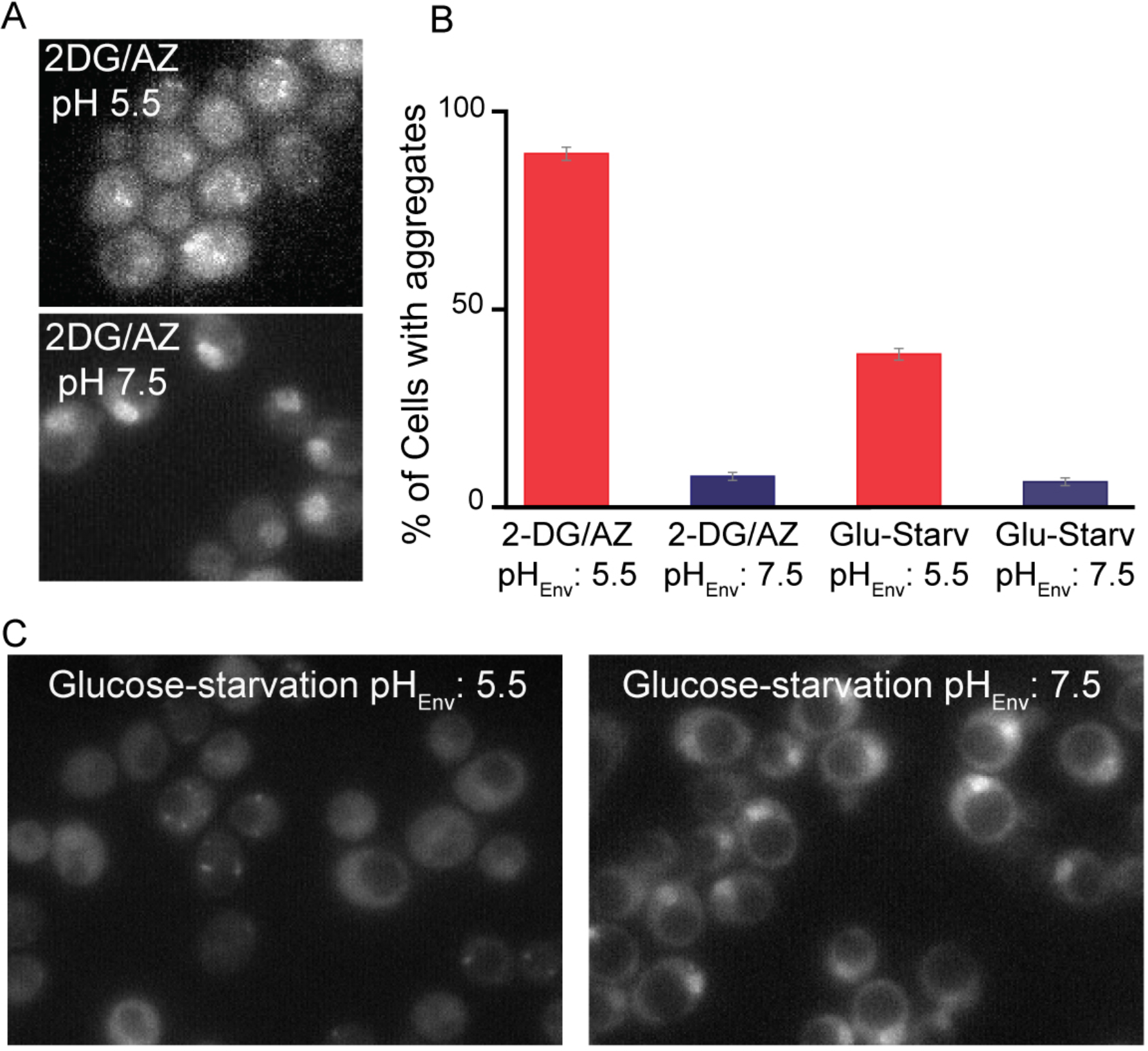
A synthetic mini-spidroin protein aggregates upon glucose-starvation at pH_env_ 5.5 but no at pH_env_ 7.5. A- Spidroin–GFP on exponentially growing cells. B- Spidroin-GFP upon glucose-starvation at pH_env_ 5.5 and pH_env_ 7.5. C- Analysis of the data presented in B, Y axis showing the percentage of cells that have aggregates.

## Material and Methods

### Cloning and yeast transformations

Yeast strains used in this study are in the S288c strain-background (derived from BY4743). The sequences of all genes in this study were obtained from the *Saccharomyces cerevisiae* genome database (http://www.yeastgenome.org/).

We cloned the various *SNF5* mutants into plasmids from the Longtine/Pringle collection (Longtine *et al*., 1998). We assembled plasmids by PCR or gene synthesis (IDT gene-blocks) followed by Gibson cloning (Gibson *et al*., 2009). Then, plasmids were linearized and used to replace the WT locus by sigma homologous recombination at both ends of the target gene.

The *ΔQ-SNF5* gene lacks the N-terminal 282 amino acids that comprise a glutamine rich low complexity domain. Methionine 283 serves as the ATG for the *ΔQ-SNF5* gene. In the *_HtoA-_snF5* allele, histidines 106, 109, 213 and 214 were replaced by alanine using mutagenic primers to amplify three fragments of the polyQ region which were combined by Gibson assembly into a *SNF5* parent plasmid linearized with BamH1 and Sac1.

We noticed that the slow growth null strain phenotype of the *snf5Δ* was partially lost over time, presumably due to suppressor mutations. Therefore, to avoid spontaneous suppressors, we first supplemented WT S288c with a CEN/ARS plasmid carrying the *SNF5* gene under its own promoter and the *URA3* auxotrophic selection marker. Then a KanMX resistance cassette, amplified with primers with homology at the 5’ and 3’ of the *SNF5* gene was used to delete the entire chromosomal *SNF5* ORF by homologous recombination. We cured strains of the CEN/ARS plasmid carrying WT *SNF5* by negative selection against its URA3 locus by streaking for single colonies on 5-FOA plates immediately before each experiment to analyze the *snf5Δ* phenotype.

The *ADH2* reporter was cloned into pRS collection plasmids (Chee and Haase, 2012) for integration. *URA3* (pRS306) or *LEU2* (pRS305) were used as auxotrophic selection markers. The 835 base pairs upstream of the +1 of the *ADH2* gene and the mCherry ORF were amplified by PCR and assembled into linearized pRS plasmids (Sac1/Asc1) by Gibson assembly. These plasmids were cut with Sph1 in the middle of the *ADH2* promoter and integrated into the endogenous *ADH2* locus by homologous recombination.

The *pHluorin* gene was clone on a pRS collection plasmid for integration. *URA3* (pRS306) and *LEU2* (pRS305) were used for selection. The plasmid with the *pHluorin* gene was obtained (Orij *et al*., 2009). We amplified the *pHluorin* gene and the strong TDH3 promoter and used Gibson assembly to clone these fragments into pRS plasmids linearized with Sac1 and Asc1.

A C-terminal TAP tag was used to visualize Snf5 and Snf2 proteins in Western blots. pRS plasmids were used but the cloning strategy was slightly different. A C-terminal region of the *SNF5* and SNF2 genes were PCR amplified without the Stop codon. This segment does not contain a promoter or an ATP codon for translation initiation. The TAP tag was then amplified by PCR and cloned together with the 3’ of *SNF5* and *SNF2* by Gibson assembly into pRS plasmids with linearized Sac1 and Asc1. The plasmids linearized in the *snf5* 3’ and snf2 3’ with StuI and XbaI respectively done to linearize the plasmid allowing integration it into the 3’ of each gene locus by homologous recombination. Therefore, after transformation the WT promoter is upstream of the WT gene without the stop codon and fused to the TAP tag.

GFP strains, the *SNF5*-*GFP* strain was obtained by the yeast GFP collection (Huh *et al*., 2003), a gift of the Drubin/Barnes laboratory at UC Berkeley. The *SNF2-GFP* fused strain was made by the same approached used for the TAP tagged strain above.

### Culture media

Most experiments, unless indicated, were performed in synthetic complete media (13.4 g/L yeast nitrogen base and ammonium sulfate; 2 g/L amino acid mix and 2% glucose). Carbon starvation media was synthetic complete without dextrose and supplemented with sorbitol, a non-fermentable carbon source to avoid osmotic shock during glucose-starvation (6.7 g/L YNB + ammonium sulfate; 2g/L Amino acid mix and 100 mM Sorbitol). pH was adjusted using 5 M NaOH.

### Glucose-starvation

Cultures were incubated in a rotating incubator at 30^Ο^C and grown overnight (14-16 h) to an OD between 0.2 and 0.3. Note: it is extremely important to prevent culture OD from exceeding 0.3 – and results are different if cells are allowed to saturate and then diluted back. Thus, it is imperative to obtain log phase cultures directly to obtain reproducible results. 3 milliliters of OD 0.2-0.3 culture were centrifuged at 6000 RPM for 3 minutes, re-suspended in SC Sorbitol at different pHs and washed 2 times. Finally, cells were re-suspended in 3 milliliters of SC Sorbitol. For flow cytometry, aliquots of 200 uL were taken for each time point in 96 well plates. During the course of time lapse experiments, culture aliquots were set aside at 4^Ο^C. LSR II – HTS were used for all measurements. 10,000 cells were measured for each time point.

### Cytosolic pH measurements

The cytosolic pH measurements were made with flow cytometry or microscopy. *pHluorin* was used to measure pH based on the ratio of fluorescence from two excitation wavelengths. In our cytometry we used the settings for AmCyan (excitation 457, emission 491) and FITC (excitation 494, emission 520). While the AmCyan emission increases with pH, FITC emission decreases. A calibration curve was made in each experiment for each strain. To generate a calibration curve, glycolysis and respiration were poisoned using 2-deoxyglucose and azide leading to equilibration of the cytosolic pH to the extracellular pH. Although many buffers are available, we used the calibration buffer published by Patricia Kane’s group (Diakov, Tarsio and Kane, 2013): 50 mM MES (2-(N-morpholino) ethanesulfonic acid), 50 mM HEPES (4-(2-hydroxyethyl)-1-piperazineethanesulfonic acid, 50 mM KCL, 50 mM NaCL, 0.2 M ammounium acetate, 10 mM sodium azide, 10 mM 2-Deoxyglucose. Buffers were titrated to pH with HCL and NaOH to the desired pH. Sodium Azide and 2-deoxyglucose was always added fresh.

### RT-Q-PCR

For qPCR and RNA seq, RNA was extracted with the “High pure RNA isolation kit” (Roche) following the manufacturer instructions. Three biological replicates were done for qPCR and RNAseq. cDNAs and qPCR were made with iSCRIPT and iTAQ universal SYBR green supermix by Bio-Rad, following the manufacturer instructions. Samples processed were: Exponentially growing culture (+Glu) and glucose-starvation at pH 5.5 and 7.5 for 4 hours. Primers qPCR were taken from Biddick et al 2008. For *ADH2* and *FBP1* genes are: Forward (GTC TAT CTC CAT TGT CGG CTC)/ Reverse (GCC CTT CTC CAT CTT TTC GTA) and Forward (CTT TCT CGG CTA GGT ATG TTG G)/ Reverse (ACC TCA GTT TTC CGT TGG G). *ACT1* was used as control amplification: Forward (TGG ATT CCG GTG ATG GTG TT)/ Reverse (TCA AAA TGG CGT GAG GTA GAG A).

### RNA sequencing

We performed RNA sequencing analysis to determine the extent of the requirement for the*SNF5* polyQ domains in the activation of glucose-repressed genes. Total RNA was extracted from WT, *ΔQ-snf5* and *_HtoA-_snf5* strains during exponentially growth (+Glu) and after 4 hours of glucose starvation. Next, Poly-A selection was performed using Dynabeads and libraries were performed following manufactures indications. Sequencing of the 32 samples was performed on an Illumina Hi-seq on two lanes. RNA-seq data were aligned to the University of California, Santa Cruz (UCSC), sacCer2 genome using Kallisto (0.43.0, http://www.nature.com/nbt/journal/v34/n5/full/nbt.3519.html) and downstream visualization and analysis was done using R (3.2.2). Differential gene expression analysis and heat maps were created using Sleuth where a least ratio test was used to determine differentially expressed genes and Euclidean distance to calculate clustering for heat maps.

### Western blots

Strains containing *SNF5* and *SNF2* fused to the TAP tag were used. Given the low concentration of these proteins, they were extracted with Trichloroacetic acid (TCA): 3 mL or a colony (re-suspended in water) were pelleted by centrifugation for 2 min at 6000 RPM and then frozen in liquid nitrogen. Pellets were thawed on ice and re-suspended in 200 uL of 20% TCA, ~0.4 g of glass beads were added to each tube. Samples were lysed by bead beating 4 times for 2 min with 2 min of resting in ice in each cycle. Supernatants were extracted using a total of 1 mL of 5% TCA and precipitated for 20 min at 14000 RPM at 4 C. Finally, pellets were re-suspended in 212 uL of Laemmli sample buffer and pH adjusted with ~26 uL of Tris buffer pH 8. Samples were run on 7-12% gradient acrylamide gels with Thermo-Fisher PageRuler Prestained protein ladder 10 to 18 KDa. Once transferred, the membrane was blocked with 5% milk and incubated with a rabbit IgG primary antibody (which is bound by the protein A moiety of the TAP tag) for 1 hour and then with goat anti-rabbit secondary antibody. Membranes were visualized using LiCor Odyssey CLx scanner with Image studio 3.1 software. Membranes were scanned at 700 and 800 nM excitation light.

### Data fitting into Gaussian curves

The *ADH2* expression inferred from a fluorescent reporter (*P_ADH2_-mCherry)* and cytosolic pH were fitted with a single or double Gaussian curve for statistical analysis, with a Matlab function developed in the lab. The choice of single or double Gaussian fit was determined by assessing which fit gave the least residuals. The height of the single or double Gaussian were used to determine the fraction of cells in each peaks. For simplicity, we quantified the height of the Gaussian rather than the area because peaks overlapped in many conditions.

### Sequence analysis of polyQs

A polyQ structure in protein sequences was defined as a polypeptide sequence containing at least ten glutamines allowing any number of single or double amino-acid insertions, but broken by any interruption of three or more non-glutamine amino acid residues. For example, QQQQQAAQQQQQ and QAQAQAQAQAQAQAQAQAQ both count as polyQ, but QQQQQAAAQQQQQ does not. *Saccharomyces cerevisiae* genome and protein sequences (S288c) were downloaded from SGD (www.yeastgenome.org). S288c transversion and transition rates were obtained from Zhu *et al*, 2014. We ran a computational evolution experiment with a single nucleotide mutation probability of 0.012 per nucleotide. This simulation leads to a non-synonymous change in roughly 20% of the glutamines, thus maintaining the overall integrity of the polyQ structure but introducing enough changes for statistical analysis. We ran this simulation ten thousand times using the polyQ-encoding portion of the S288c reference genome. Additionally, ten thousand simulations were performed on artificial genomes consisting of pure ‘CAA’, ‘CAG’ or ‘CAGCAA’ until we obtained the same non-glutamine frequency within the polyQ structure as found in S288c protein sequences. We defined an enrichment score as the relative frequency of a specific amino acid residue within or surrounding a polyQ normalized by the overall frequency of this amino acid residue present in the whole reference genome. Besides *Saccharomyces cerevisiae*, enrichment scores within and around polyQ were calculated for *Drosophila melanogaster*, *Homo sapiens* and *Dictyostelium discoideum* reference protein sequences (downloaded from http://www.ebi.ac.uk) (Zhu *et al*., 2014).

## References

Abrams, E., Neigeborn, L. and Carlson, M. (1986) ‘Molecular analysis of SNF2 and SNF5, genes required for expression of glucose-repressible genes in Saccharomyces cerevisiae.’, Molecular and cellular biology, 6(11), pp. 3643–51. doi: 10.1128/mcb.6.11.3643.

Askarieh, G., Hedhammar, M., Nordling, K., Saenz, A., Casals, C., Rising, A., Johansson, J. and Knight, S. D. (2011) ‘Self-assembly of spider silk proteins is controlled by a pH-sensitive relay’, Nature. Nature Publishing Group, 465(7295), pp. 236–238. doi: 10.1038/nature08962.

Bates, G. P., Dorsey, R., Gusella, J. F., Hayden, M. R., Kay, C., Leavitt, B. R., Nance, M., Ross, C. A., Scahill, R. I., Wetzel, R., Wild, E. J. and Tabrizi, S. J. (2015) ‘Huntington disease’, Nature Reviews Disease Primers, (April), p. 15005. doi: 10.1038/nrdp.2015.5.

Biddick, R. K., Law, G. L., Chin, K. K. B. and Young, E. T. (2008) ‘The transcriptional coactivators SAGA, SWI/SNF, and mediator make distinct contributions to activation of glucose-repressed genes’, Journal of Biological Chemistry, 283(48), pp. 33101–33109. doi: 10.1074/jbc.M805258200.

Biddick, R. K., Law, G. L. and Young, E. T. (2008) ‘Adr1 and Cat8 mediate coactivator recruitment and chromatin remodeling at glucose-regulated genes’, PLoS ONE, 3(1). doi: 10.1371/journal.pone.0001436.

Biegel, J. A., Zhou, J., Rorke, L. B., Stenstrom, C., Wainwright, L. M. and Fogelgren, B. (1999) ‘Advances in Brief Germ-Line and Acquired Mutations of INI1 in Atypical Teratoid and Rhabdoid Tumors’, pp. 74–79.

Brangwynne, C. P., Mitchison, T. J. and Hyman, A. A. (2011) ‘Active liquid-like behavior of nucleoli determines their size and shape in Xenopus laevis oocytes’, Proc. Natl. Acad. Sci., 108(11), pp. 4334–4339. doi: 10.1073/pnas.1017150108.

Busa, W. B. and Crowe, J. H. (1983) ‘Intracellular pH Regulates Transitions between Dormancy and Development of Brine Shrimp (Artemia salina) Embryos’, Science, 221(4608), pp. 366–368. doi: 10.1126/science.221.4608.366.

Busa, W. B. and Nuccitelli, R. C. N.-C. (1984) ‘Metabolic regulation via intracellular pH’, Am J Physiol Regul Integr Comp Physiol, 246, pp. R409–R438.

Carlson, M. (1987) ‘Regulation of sugar utilization in Saccharomyces species.’, Journal of Bacteriology, 169(11), pp. 4873–4877.

Chee, M. K. and Haase, S. B. (2012) ‘New and Redesigned pRS Plasmid Shuttle Vectors for Genetic Manipulation of Saccharomyces cerevisiae’, G3&#58; Genes|Genomes|Genetics, 2(5), pp. 515–526. doi: 10.1534/g3.111.001917.

Chiba, H., Muramatsu, M., Nomoto, A. and Kato, H. (1994) ‘Two human homologues of saccharomyces cerevisiae SWI2/SNF2 and Drosophila brahma are transcriptional coactivators cooperating with the estrogen receptor and the retinoic acid receptor’, Nucleic Acids Research, 22(10), pp. 1815–1820. doi: 10.1093/nar/22.10.1815.

DeRisi, J. L. (1997) ‘Exploring the Metabolic and Genetic Control of Gene Expression on a Genomic Scale’, Science, 278(5338), pp. 680–686. doi: 10.1126/science.278.5338.680.

Diakov, T. T., Tarsio, M. and Kane, P. M. (2013) ‘Measurement of vacuolar and cytosolic pH in vivo in yeast cell suspensions.’, Journal of visualized experiments : JoVE, (74), pp. 1–7. doi: 10.3791/50261.

Dutta, A., Sardiu, M., Gogol, M., Gilmore, J., Zhang, D., Florens, L., Abmayr, S. M., Washburn, M. P. and Workman, J. L. (2017) ‘Composition and Function of Mutant Swi/Snf Complexes’, Cell Reports. ElsevierCompany., 18(9), pp. 2124–2134. doi: 10.1016/j.celrep.2017.01.058.

Fan, H., Ho, L., Chi, C., Chen, S., Peng, G., Chan, T., Lin, S. and Harn, H. (2014) ‘Review Polyglutamine (PolyQ) Diseases : Genetics to Treatments’, 23(235), pp. 441–458. doi: 10.3727/096368914X678454.

Gagliardi, L. J. and Shain, D. H. (2013) ‘Is intracellular pH a clock for mitosis?’, Theoretical biology & medical modelling, 10(1), p. 8. doi: 10.1186/1742-4682-10-8.

Geng, F., Cao, Y. and Laurent, B. C. (2001) ‘Essential Roles of Snf5p in Snf-Swi Chromatin Remodeling In Vivo Essential Roles of Snf5p in Snf-Swi Chromatin Remodeling In Vivo’, Society, 21(13), pp. 4311–4320. doi: 10.1128/MCB.21.13.4311.

Gibson, D. G., Young, L., Chuang, R.-Y., Venter, J. C., Hutchison, C. a, Smith, H. O., Iii, C. A. H. and America, N. (2009) ‘Enzymatic assembly of DNA molecules up to several hundred kilobases.’, Nature methods, 6(5), pp. 343–5. doi: 10.1038/nmeth.1318.

Hafke, J. B., Neff, R., Hütt, M. T., Lüttge, U. and Thiel, G. (2001) ‘Day-to-night variations of cytoplasmic pH in a crassulacean acid metabolism plant.’, Protoplasma, 216(3–4), pp. 164–70.

Huh, W.-K., Falvo, J. V., Gerke, L. C., Carroll, A. S., Howson, R. W., Weissman, J.S. and O’Shea, E. K. (2003) ‘Global analysis of protein localization in budding yeast’, Nature, 425(6959), pp. 686–691. doi: 10.1038/nature02026.

Huntley, M. A. and Clark, A. G. (2007) ‘Evolutionary analysis of amino acid repeats across the genomes of 12 drosophila species’, Molecular Biology and Evolution, 24(12), pp. 2598–2609. doi: 10.1093/molbev/msm129.

Janody, F., Sturny, R., Schaeffer, V., Azou, Y. and Dostatni, N. (2001) ‘Two distinct domains of Bicoid mediate its transcriptional downregulation by the Torso pathway.’, Development (Cambridge, England), 128(12), pp. 2281–90. doi: 10.1038/374657a0.

Joyner, R. P., Tang, J. H., Helenius, J., Dultz, E., Brune, C., Holt, L. J., Huet, S., M??ller, D. J. and Weis, K. (2016) ‘A glucose-starvation response regulates the diffusion of macromolecules’, eLife, 5(MARCH2016), pp. 1–26. doi: 10.7554/eLife.09376.

Kadoch, C., Hargreaves, D. C., Hodges, C., Elias, L., Ho, L., Ranish, J. and Crabtree, G. R. (2013) ‘a n a ly s i s Proteomic and bioinformatic analysis of mammalian SWI / SNF complexes identifies extensive roles in human malignancy’, Nature Publishing Group. Nature Publishing Group, 45(6), pp. 592–601. doi: 10.1038/ng.2628.

Kadonaga, J. T., Carner, K. R., Masiarz, F. R. and Tjian, R. (1987) ‘Isolation of cDNA encoding transcription factor Sp1 and functional analysis of the DNA binding domain.’, Cell, 51(6), pp. 1079–1090. doi: 0092-8674(87)90594-0 [pii].

Kadonaga, J. T., Courey, A. J., Ladika, J. and Tjian, R. (1988) ‘Distinct regions of Sp1 modulate DNA binding and transcriptional activation’, Science, 242(4885), pp. 1566–1570.

Kane, P. M. (1995) ‘Disassembly and Reassembly of the Yeast Vacoular H+ ATPase in vivo’, The Journal of Biological Chemistry, pp. 17025–17032. doi: 10.1074/jbc.270.28.17025.

Karagiannis, J. and Young, P. G. (2001) ‘Intracellular pH homeostasis during cell-cycle progression and growth state transition in Schizosaccharomyces pombe.’, Journal of cell science, 114(Pt 16), pp. 2929–41. doi: 10.1091/mbc.E04.

Kato, M., Han, T. W., Xie, S., Shi, K., Du, X., Wu, L. C., Mirzaei, H., Goldsmith, E. J., Longgood, J., Pei, J., Grishin, N. V., Frantz, D. E., Schneider, J. W., Chen, S., Li, L., Sawaya, M. R., Eisenberg, D., Tycko, R. and McKnight, S. L. (2012) ‘Cell-free Formation of RNA Granules: Low Complexity Sequence Domains Form Dynamic Fibers within Hydrogels’, Cell, 149(4), pp. 753–767. doi: 10.1016/j.cell.2012.04.017.

Kuiper, E. F. E., de Mattos, E. P., Jardim, L. B., Kampinga, H. H. and Bergink, S. (2017) ‘Chaperones in Polyglutamine Aggregation: Beyond the Q-Stretch’, Frontiers in Neuroscience, 11(March), pp. 1–11. doi: 10.3389/fnins.2017.00145.

Kwon, I., Kato, M., Xiang, S., Wu, L., Theodoropoulos, P., Mirzaei, H., Han, T., Xie, S., Corden, J. L. and McKnight, S. L. (2013) ‘Phosphorylation-Regulated Binding of RNA Polymerase II to Fibrous Polymers of Low-Complexity Domains’, Cell. Elsevier, 155(5), pp. 1049–1060. doi: 10.1016/j.cell.2013.10.033.

Laurent, B. C., Treitel, M. a and Carlson, M. (1990) ‘The SNF5 protein of Saccharomyces cerevisiae is a glutamine- and proline-rich transcriptional activator that affects expression of a broad spectrum of genes.’, Molecular and cellular biology, 10(11), pp. 5616–25. doi: 10.1128/MCB.10.11.5616.Updated.

Levy, S. F., Ziv, N. and Siegal, M. L. (2012) ‘Bet Hedging in Yeast by Heterogeneous, Age-Correlated Expression of a Stress Protectant’, 10(5). doi: 10.1371/journal.pbio.1001325.

Li, L., Liu, H., Dong, P., Li, D., Legant, W. R., Grimm, J. B., Lavis, L. D., Betzig, E., Tjian, R. and Liu, Z. (2016) ‘Real-time imaging of Huntingtin aggregates diverting target search and gene transcription’, eLife, 5(AUGUST), pp. 1–29. doi: 10.7554/eLife.17056.001.

Longtine, M. S., McKenzie, A., Demarini, D. J., Shah, N. G., Wach, A., Brachat, A., Philippsen, P. and Pringle, J. R. (1998) ‘Additional modules for versatile and economical PCR-based gene deletion and modification in Saccharomyces cerevisiae’, Yeast, 14(10), pp. 953–961. doi: 10.1002/(SICI)1097-0061(199807)14:10<953::AIDYEA293>3.0.CO;2-U.

Martínez-Muñoz, G. A. and Kane, P. (2008) ‘Vacuolar and plasma membrane proton pumps collaborate to achieve cytosolic pH homeostasis in yeast’, Journal of Biological Chemistry, 283(29), pp. 20309–20319. doi: 10.1074/jbc.M710470200.

Miesenböck, G., De Angelis, D. a and Rothman, J. E. (1998) ‘Visualizing secretion and synaptic transmission with pH-sensitive green fluorescent proteins.’, Nature, 394(6689), pp. 192–5. doi: 10.1038/28190.

Munder, M. C., Midtvedt, D., Franzmann, T., Nüske, E., Otto, O., Herbig, M., Ulbricht, E., Müller, P., Taubenberger, A., Maharana, S., Malinovska, L., Richter, D., Guck, J., Zaburdaev, V. and Alberti, S. (2016) ‘A pH-driven transition of the cytoplasm from a fluid- to a solid-like state promotes entry into dormancy’, eLife. eLife Sciences Publications Limited, 5, pp. 59–69. doi: 10.7554/eLife.09347.

Needham, J. (1926) ‘The Hydrogen-Ion Concentration and Oxidation-Reduction Potential of the Cell-Interior before and after Fertilisation and Cleavage : A Micro-Injection Study on Marine Eggs Author (s): Joseph Needham and Dorothy Moyle Needham Source : Proceedings of the R’, 99(695), pp. 173–199.

Neigeborn, L. and Carlson, M. (1984) ‘GENES AFFECTING THE REGULATION OF SUC2 GENE EXPRESSION BY GLUCOSE REPRESSION IN SACCHAROMYCES CEREVISIAE’, Genetics, 108, pp. 845–858.

Nickerson, J. A., Wu, Q., Imbalzano, A. N. and Nickerson, J. A. (2017) ‘Mammalian Swi / SnF enzymes and the epigenetics of Tumor Cell Metabolic Reprogramming’, 7(April), pp. 1–9. doi: 10.3389/fonc.2017.00049.

Okamoto, Y. K. K. (1994) ‘Cytoplasmic Ca+2 and H+ concentrations determine cell fate in Dictyostelium discoideum’, Access, 28(13), pp. 2423–2427.

Orij, R., Postmus, J., Beek, A. Ter, Brul, S. and Smits, G. J. (2009) ‘In vivo measurement of cytosolic and mitochondrial pH using a pH-sensitive GFP derivative in Saccharomyces cerevisiae reveals a relation between intracellular pH and growth’, Microbiology, 155(1), pp. 268–278. doi: 10.1099/mic.0.022038-0.

Perutz, M. F., Pope, B. J., Owen, D., Wanker, E. E. and Scherzinger, E. (2002) ‘Aggregation of proteins with expanded glutamine and alanine repeats of the glutamine-rich and asparagine-rich domains of Sup35 and of the amyloid β-peptide of amyloid plaques’, Proceedings of the National Academy of Sciences of the United States of America, 99(8), pp. 5596–5600. doi: 10.1073/pnas.042681599.

Peterson, C. L., Dingwall, A. and Scott, M. P. (1994) ‘Five SWI/SNF gene products are components of a large multisubunit complex required for transcriptional enhancement.’, Proceedings of the National Academy of Sciences of the United States of America, 91(8), pp. 2905–8.

Peterson, C. L. and Herskowitz, I. (1992) ‘Characterization of the yeast SWI1, SWI2, and SWI3 genes, which encode a global activator of transcription’, Cell, 68(3), pp. 573–583. doi: 10.1016/0092-8674(92)90192-F.

Petrovska, I., Nüske, E., Munder, M. C., Kulasegaran, G., Malinovska, L., Kroschwald, S., Richter, D., Fahmy, K., Gibson, K., Verbavatz, J. M. and Alberti, S. (2014) ‘Filament formation by metabolic enzymes is a specific adaptation to an advanced state of cellular starvation’, eLife, 2014(3), pp. 1–19. doi: 10.7554/eLife.02409.

Prochasson, P., Neely, K. E., Hassan, A. H., Li, B. and Workman, J. L. (2003) ‘Targeting activity is required for SWI/SNF function in vivo and is accomplished through two partially redundant activator-interaction domains’, Molecular Cell, 12(4), pp. 983–990. doi: 10.1016/S1097-2765(03)00366-6.

Schaefer, M. H., Wanker, E. E. and Andrade-Navarro, M. A. (2012) ‘Evolution and function of CAG/polyglutamine repeats in protein-protein interaction networks’, Nucleic Acids Research, 40(10), pp. 4273–4287. doi: 10.1093/nar/gks011.

van Schalkwyk, D. A., Saliba, K. J., Biagini, G. A., Bray, P. G. and Kirk, K. (2013) ‘Loss of pH Control in Plasmodium falciparum Parasites Subjected to Oxidative Stress’, PLoS ONE, 8(3). doi: 10.1371/journal.pone.0058933.

Sen, P., Luo, J., Hada, A., Paul, S., Ranish, J., Bartholomew, B., Hailu, S. G., Dechassa, M. L., Persinger, J. and Brahma, S. (2017) ‘Loss of Snf5 Induces Formation of an Aberrant SWI/SNF Complex Loss of Snf5 Induces Formation of an Aberrant SWI/SNF Complex’, Cell Reports. ElsevierCompany., 18(9), pp. 2135–2147. doi: 10.1016/j.celrep.2017.02.017.

Sevenet, N., Sheridan, E., Amram, D., Schneider, P., Handgretinger, R. and Delattre, O. (1999) ‘Constitutional Mutations of the hSNF5 / INI1 Gene Predispose to a Variety of Cancers’, pp. 1342–1348.

Shen, K., Calamini, B., Fauerbach, J. A., Ma, B., Shahmoradian, S. H., Serrano Lachapel, I. L., Chiu, W., Lo, D. C. and Frydman, J. (2016) ‘Control of the structural landscape and neuronal proteotoxicity of mutant Huntingtin by domains flanking the polyQ tract.’, eLife, 5, pp. 1–29. doi: 10.7554/eLife.18065.

Sudarsanam, P., Iyer, V. R., Brown, P. O. and Winston, F. (2000) ‘Whole-genome expression analysis of snf/swi mutants of Saccharomyces cerevisiae.’, Proceedings of the National Academy of Sciences of the United States of America, 97(7), pp. 3364–3369. doi: 10.1073/pnas.97.7.3364.

Tannock, I. F. and Rotin, D. (1989) ‘Perspectivesin CancerResearch Acid pH in Tumors and Its Potential for Therapeutic Exploitation1’, 9.

Thakur, A. K., Jayaraman, M., Mishra, R., Thakur, M., Chellgren, V. M., Byeon, I.-J. L., Anjum, D. H., Kodali, R., Creamer, T. P., Conway, J. F., Gronenborn, A. M. and Wetzel, R. (2015) ‘Polyglutamine disruption of the huntingtin exon 1 N terminus triggers a complex aggregation mechanism’. doi: 10.1038/nsmb.1570.

Vercoulen, Y., Kondo, Y., Iwig, J. S., Janssen, A. B., White, K. A., Amini, M., Barber, D. L., Kuriyan, J. and Roose, J. P. (2017) ‘A Histidine pH sensor regulates activation of the Ras-specific guanine nucleotide exchange factor RasGRP1’, pp. 1–26.

Whitten, S. T., Garcia-Moreno E., B. and Hilser, V. J. (2005) ‘Local conformational fluctuations can modulate the coupling between proton binding and global structural transitions in proteins’, Proceedings of the National Academy of Sciences, 102(12), pp. 4282–4287. doi: 10.1073/pnas.0407499102.

Wike-Hooley, J. L., J. Haveman, J. and Reinhold, H. S. (1984) ‘The relevance of tumour pH to the treatment of malignant disease’, 2, pp. 343–366.

Wu, Q., Madany, P., Dobson, J. R., Schnabl, J. M., Sharma, S., Smith, T. C., Wijnen, A. J. Van, Stein, J. L., Lian, J. B., Stein, G. S., Muthuswami, R., Imbalzano, A. N. and Nickerson, J. A. (2016) ‘The BRG1 chromatin remodeling enzyme links cancer cell metabolism and proliferation’, 7(25).

Young, B. P., Shin, J. J. H., Orij, R., Chao, J. T., Li, S. C., Guan, X. L., Khong, A., Jan, E., Wenk, M. R., Prinz, W. A., Smits, G. J. and Loewen, C. J. R. (2010) ‘Phosphatidic Acid Is a pH Biosensor That Links Membrane Biogenesis to Metabolism’, Science, 329(5995), pp. 1085–1088. doi: 10.1126/science.1191026.

Young, E. T., Tachibana, C., Chang, H.-W. E., Dombek, K. M., Arms, E. M. and Biddick, R. (2008) ‘Artificial recruitment of mediator by the DNA-binding domain of Adr1 overcomes glucose repression of ADH2 expression’, Molecular and cellular biology, 28(8), pp. 2509–2516. doi: 10.1128/MCB.00658-07.

Zhang, H., Elbaum-Garfinkle, S., Langdon, E. M., Taylor, N., Occhipinti, P., Bridges, A. A., Brangwynne, C. P. and Gladfelter, A. S. (2015) ‘RNA Controls PolyQ Protein Phase Transitions’, Molecular Cell. Elsevier Inc., 60(2), pp. 220–230. doi: 10.1016/j.molcel.2015.09.017.

Zhu, Y. O., Siegal, M. L., Hall, D. W. and Petrov, D. A. (2014) ‘Precise estimates of mutation rate and spectrum in yeast.’, Proceedings of the National Academy of Sciences of the United States of America, 111(22), pp. E2310-8. doi: 10.1073/pnas.1323011111.

Zid, B. M. and O’Shea, E. K. (2014) ‘Promoter sequences direct cytoplasmic localization and translation of mRNAs during starvation in yeast. TL - 514’, Nature. Nature Publishing Group, 514 VN-(7520), pp. 117–121. doi: 10.1038/nature13578.

